# The CNS lymphatic system modulates the adaptive neuro-immune response in the perilesional cortex in a mouse model of traumatic brain injury

**DOI:** 10.1101/821645

**Authors:** Wojciechowski Sara, Vihma Maria, Galbardi Barbara, Virenque Anaïs, Meike H. Keuters, Antila Salli, Alitalo Kari, Koistinaho Jari, Francesco M. Noe

**Affiliations:** A.I. Virtanen Institute for Molecular Sciences, University of Eastern Finland, 70210 Kuopio, Finland; Neuroscience Center, Helsinki Institute of Life Science (HiLIFE), University of Helsinki, 00290 Helsinki, Finland; Breast Cancer Unit, Department of Medical Oncology, IRCCS Ospedale San Raffaele, 20132 Milano, Italy; Wihuri Research Institute and Translational Cancer Medicine Program, Biomedicum Helsinki, University of Helsinki, 00290 Helsinki, Finland

**Author notes:** **Correspondence:** Dr. Francesco M. Noé, HiLIFE – Neuroscience Center, University of Helsinki, Helsinki, Finland.

**Keywords:** Controlled Cortical Impact (CCI), meningeal lymphatic vessels, CD8+ T lymphocytes, resident memory T cells, deep cervical lymph nodes

## Abstract

**Rationale:** The recently discovered meningeal lymphatic vessels (mLVs) have been proposed to be the missing link between the immune and the central nervous systems. The role of mLVs in modulating the neuro-immune response following a brain injury, however, has not been analyzed. Parenchymal T lymphocyte infiltration has been previously reported as part of secondary events after traumatic brain injury (TBI), suggestive of an adaptive neuro-immune response. The phenotype of these cells has remained mostly uncharacterized. In this study, we identified the subpopulations of T cells infiltrating the perilesional areas 30 days post-injury (an early-chronic time point). Furthermore, we analyzed how the lack of mLVs affects the magnitude and the type of immune response in the brain after TBI.

**Methods:** TBI was induced in K14-VEGFR3-Ig transgenic (TG) mice or in their littermate controls (WT; wild type), applying a controlled cortical impact (CCI). One month after TBI, T cells were isolated from cortical areas ipsilateral or contralateral to the trauma and from the spleen, then characterized by flow cytometry. Lesion size in each animal was evaluated by MRI.

**Results:** In both WT and TG-CCI mice, we found a prominent T cell infiltration in the brain confined to the perilesional cortex and hippocampus. The majority of infiltrating T cells were cytotoxic CD8+ expressing a CD44^hi^CD69+ phenotype, suggesting that these are effector resident memory T cells. K14-VEGFR3-Ig mice showed a significant reduction of infiltrating CD4+ T lymphocytes, implying that mLVs are important in establishing a proper neuro-immune response. Extension of the lesion (measured as lesion volume from MRI) did not differ between the genotypes. Finally, TBI did not relate with alterations in peripheral circulating T cells, as assessed one month after injury induction.

**Conclusions:** Our data support the hypothesis that mLVs are pivotal for a proper and specific neuro-immune response after TBI, which is principally mediated by the resident memory CD8+ T cells.

## Introduction

Traumatic brain injury (TBI) is among the top causes of death and disability in adult life. (Hale et al., 2019, Hyder et al., 2007). At least 70 million people worldwide are estimated to incur TBIs every year (Dewan et al., 2018), with the number of prevalent cases of TBI in 2016 above 55 million, suffering from a wide range of lifelong physical and psychological invalidities (GBD 2016 Neurology Collaborators, 2019).

TBI is defined as an alteration in brain function, or other evidence of brain pathology, caused by an external force (Menon et al., 2010), which results in immediate neuronal cell death, diffuse axonal injury, ischemia, and hemorrhage (McIntosh et al., 1996). These primary insults initiate a progressive cascade of secondary injuries, which include macrophage infiltration (Braun et al., 2017), neuro-inflammation (microglia and astrocyte activation associated with cytokine production), edema formation, oxidative stress, neuronal necrosis and apoptosis, and white matter atrophy (McIntosh et al., 1996). Secondary injuries can progress for years in patients and rodent models of TBI, and are the causes of the neurological and psychiatric deficits associated with the pathology (DeKosky et al., 1998).

Among secondary events following TBI, recruitment of peripheral immune cells into the brain, including T lymphocytes, has been described (Daglas et al., 2019, Bai et al., 2017, Erturk et al., 2016, Jin et al., 2012). Two distinct waves of infiltrating CD3+ T cells have been reported in the injured brain. First, a massive infiltration immediately commences after trauma and peaks 3 days post-injury (dpi) (Jin et al., 2012). After one month, there is a late adaptive immune response with a second recruitment, which persists chronically (Erturk et al., 2016). However, the mechanisms and the consequences of the activation of the adaptive immune system after TBI are still poorly understood.

A proper immune surveillance of the brain was long disputed (Galea et al., 2007), due to the lack of a classical lymphatic system within the central nervous system (CNS). However, recent studies have described the presence of anatomically distinct lymphatic vessels in the meninges surrounding the brain and the spinal cord. These meningeal lymphatic vessels (mLVs) preferentially drain the cerebrospinal fluid into the deep cervical lymph nodes (dcLNs) (Louveau et al., 2018, Louveau et al., 2015, Aspelund et al., 2015). Within these secondary lymphoid organs, brain-derived antigens are presented to resident T lymphocytes, evoking different cellular fate and immune responses based on the inflammatory milieu. It has been demonstrated that dcLNs play a specific role in neuro-immune interaction, ensuring the protection of brain cells by promoting a non-cytotoxic immune response (Thomas, D. L. et al., 2008, Harling-Berg et al., 1999, Cserr et al., 1992). From this prospective, mLVs and dcLNs are essential components of a putative specific CNS lymphatic system, and the mLVs could be essential in the activation of immune responses to brain insults, by transporting brain-derived antigens to the dcLNs.

The aim of our work is to better characterize the late adaptive immune response and to decipher the mechanisms underpinning the activation of T lymphocytes after TBI, focusing on the specific role of mLVs in this process. In this regard, we induced a cerebral contusion in the cortex of transgenic K14-VEGFR3-Ig (TG) mice that completely lack lymphatic vessels in several tissues, including the meninges (Aspelund et al., 2015, Makinen et al., 2001). We examined the phenotype of T lymphocytes infiltrating the perilesional cortical areas, determining the prevalence of a CD8+ mediated cytotoxic immune response in the TBI mice lacking the lymphatic system. One month after brain injury, infiltrating T lymphocytes and circulating peripheral T cell populations in the spleen were phenotyped by flow cytometry. MRI was used to evaluate and compare lesion size in both transgenic animals and in their wild type (WT) littermates.

Our data show that the CNS immune response after TBI is specific and independent from peripheral immune activation. We also demonstrate that the lack of a functional mLVs-dcLNs connection alters the neuro-immune interaction after TBI, specifically dampening the CD4+ mediated immune response. No differences in MRI cortical lesion were found between the two genotypes. Finally, independent of genotype, infiltrating T cells present a resident memory effector phenotype, supporting their role in secondary injuries after TBI.

## Material and Methods

### Mice

Initial breeding pairs of K14-VEGFR3-Ig mice (C57BL/6JOlaHsd background (Makinen et al., 2001)) were transferred from the University of Helsinki, and the colony was further expanded and maintained at University of Eastern Finland (Kuopio, Finland). Wild type and transgenic K14-VEGFR3-Ig mice used in all the experiments were littermates. Genotype screening was routinely confirmed by polymerase chain reaction analysis of ear punch samples. Mixed WT and TG mice were housed in standard laboratory cages (four animals per cage, until surgery) in a controlled enriched environment (constant temperature, 22□±□1□°C, humidity 50–60 %, lights on 07:00– 19:00), with food and water available *ad libitum* (Hutchinson et al., 2005). After TBI induction, mice were kept two per cage, separated individually by a pierced partition. All animal procedures were approved by the Animal Ethics Committee of the Provincial Government of Southern Finland (ESAVI-2018-008787) and performed in accordance with the guidelines of the European Community Council Directives 2010/63/EU.

### Controlled cortical Injury (CCI) mouse model of TBI

All surgical procedures were performed aseptically whenever possible. Adult, 5 month-old male mice were deeply anesthetized with isoflurane (5 % for induction, 1.0-1.5 % for maintenance, in 0.5 L/min air; see Supplementary Table 1), injected with Carprofen (4 mg/Kg; s.c.) and the heads fixed to a stereotaxic frame (Kopf, Tujunga, USA). The scalp was shaved and then scrubbed (3x) with alternating Betadine (povidone-iodine 10 %) and 70 % ethanol, then local anesthesia of 2% Xylocain gel was applied. After skull exposure, a 5 mm circular craniotomy was manually drilled over the left parieto-temporal cortex, with the posterior edge of the craniotomy opposed to the lambdoid suture and the right edge to the sagittal suture. In order to reduce heating during manual craniotomy, the skull was irrigated with cold 0.9 % saline solution. The carved bone was carefully removed, without disrupting the underlying dura, and placed in 1 % Betadine solution. Thereafter, the animal was disconnected from isoflurane anesthesia for 5 min (stage 3 plane 1 according to Guedel’s classification (Guedel, 1927)), and CCI was induced using an electromagnetic stereotaxic impact actuator (ImpactOne, Leica, Richmond, VA, USA). The 3 mm blunt tip of the impactor was adjusted to the center of the exposed dura perpendicular to the brain surface, and the impact was administered at a depth of 0.5 mm, speed of 5.0 m/s, and dwell time of 100 ms. The total duration of the craniotomy procedure including anesthesia induction was 35-40 min (Supplementary Table 1). After the impact, the mouse was reconnected to the isoflurane system and the skull secured with bone cement (Selectaplus + Palacos R+G 50/50). The scalp was sutured and treated with Cicatrene powder (Neomycin + Bacitracin) and Terramycin spray (Oxytetracycline). The total duration of post-impact surgery was 10 min. The mice were injected i.p. with 1 mL pre-warmed sterile saline (35 °C) and allowed to fully recover in an incubator at 32 °C. Mice were followed for the subsequent 48 h for any signs of illness or distress, in which case Carprofen was administered. Daily examination was performed for general health/mortality, and moribundity for the rest of the study. No mortality was observed.

Craniotomy-related neuroinflammation has been previously reported in this model and the craniotomy itself can be considered a form of mild brain trauma (Sashindranath et al., 2015, Cole et al., 2011). Moreover, CCI is a model of penetrating injury, involving dura damage, which has a severity that bypasses the possible effect of meningeal inflammation related to the craniotomy. The aim of our study is to characterize the adaptive immunity in response to a moderate TBI. It is not to analyze how differences in trauma severity can affect the neuro-immune response. In compliance to the 3R principle, we excluded the sham-operated animals and used naïve mice not exposed to the surgical procedure as proper controls.

### In vivo MRI and lesion volume definition

MRI data were acquired 21 days after TBI induction in a 7T horizontal magnet (Bruker Pharmascan, Ettlingen, Germany). Images were acquired using a four-channel mouse brain surface coil, a 3D T2-weighted Fast Spin-Echo sequence (RARE, repetition time 1.5 s, effective echo time 48 ms, 16 echoes per excitation) with 100 μm isotropic resolution (field of view 25.6 mm x 128.8 mm x 9.6 mm; acquisition matrix 128 x 256 x 96). Scans were performed with the mouse under 1.0-1.5 % maintenance isoflurane anesthesia (70/30 N_2_O/oxygen gas mixture, 1 L/min). The average acquisition time was 40 min, including anesthesia induction. A pressure sensor was used to monitor the respiratory rate, and respiratory gating was used to minimize motion artifacts.

T2-weighted images were used to evaluate the extent of the lesion (Figure 6 and Supplementary Figure 4). Regions of interest (ROIs) were outlined for volumetric analysis, avoiding the brain-skull interface and ventricles, throughout the entire extension of the brain (excluding olfactory bulbs and cerebellum). Lesion was defined as cortical/subcortical areas with hyper-intense signal (cystic lesion) and/or signal void areas (tissue cavity) from T2-weighted images (Immonen, R. et al., 2010, Immonen, R. J., Kharatishvili, Niskanen et al., 2009). Volumes of the lesion and of the ipsilateral and contralateral hemispheres were measured using Aedes (http://aedes.uef.fi), an in-house written MatLab program (MathWorks, Natick, MA). The lesion volume and the volumes of ipsilateral and contralateral healthy hemispheres were calculated from 80 consecutive slices in the coronal plane and adjusted in the sagittal plane (66 slices) and in the axial plane (99 slices) with a volume resolution of 200 x 500 x 100 μm.

### Quantification of brain contusion area and brain atrophy

Measured volumes from MRI analysis were used to quantify the volume of the brain contusion and the brain atrophy, as previously described (Dhungana et al., 2013, Shuaib et al., 2002). The relative percentage of infarct volume was calculated using the following formula: contusion volume = (volume of contralateral hemisphere – (volume of ipsilateral hemisphere – measured lesion volume))/volume of contralateral hemisphere. Brain atrophy was determined with the following formula: atrophy = (volume of ipsilateral hemisphere – volume of contralateral hemisphere)/volume of contralateral hemisphere.

Analysis was performed blinded to the study groups. The contusion volume was measured from 22 TBI mice from the following experimental groups: wild type (WT)–CCI, n = 13; and K14-VEGFR3-Ig (TG)–CCI, n = 9.

### Cell isolation of leukocytes

Thirty days after TBI induction, mice were anesthetized with an overdose of Avertin (Sigma, St. Louis, MO, USA) then transcardially perfused with ice-cold heparinized saline. Brains were collected and placed on ice in calcium and magnesium-free Hanks Balanced salt solution (HBSS) with 25 mM HEPES (both from Sigma).

Based on the analysis of MRI, we defined *a priori* the mean extension of the lesion and of the perilesional areas for all the TBI mice. Brains were sliced using a 1 mm scored matrix (Zivic Instruments, Pittsburgh, PA, USA): 6 mm thick coronal cut encompassing the lesion area was split along the central sagittal axis into left injured and right uninjured sides. Cortical areas enclosed between the rhinal and the sagittal sulci, and the corresponding hippocampi, were further isolated, pooled together, and placed in HBSS+HEPES. From the injured sides, penetrated cortical areas were visually identified (lesion area - Supplementary Figure 1) and carefully excised along the lesion ridge to pick only the perilesional cortex for further purification of leukocytes.

Brain samples were minced with scissors and then incubated at 37 °C on a roller for 30 min in digest buffer containing 1.25 mg/mL Collagenase Type 4 (Worthington, Lakewood, NJ, USA) and 100 U/mL DNAseI (Sigma) in DMEM with GlutaMAX (Gibco Thermo Fisher Scientific, Waltham, MA, USA). Samples were filtered through a 100 μm cell strainer (Corning, Weisbaden, Germany), and centrifuged at 600 x g for 5 min. Myelin debris was removed using Debris Removal Solution (Miltenyi Biotech, Bergisch Gladbach, Germany) according to the manufacturer’s protocol. Briefly, cells were resuspended in ice-cold Dulbecco’s phosphate buffered saline (D-PBS, Sigma) with Debris Removal Solution, then overlaid with ice-cold D-PBS and centrifuged at 2500 x g for 10 min at 4 °C. Supernatant including myelin layer was carefully removed leaving the clear phase and the pellet. Samples were washed in ice-cold D-PBS, centrifuged at 600 x g for 10 min at 4 °C, and the recovered pellets were stained directly for flow cytometry.

Spleens and dcLNs were separately collected in ice-cold HBSS+HEPES and each processed by crushing through a 70 μm cell strainer (Corning). dcLNS were washed with ice-cold D-PBS containing 1% bovine serum albumin (BSA) and 2 mM ethylenediaminetetraacetic acid (EDTA), centrifuged 500 x g for 10 min and resuspended in RPMI-1640 (all from Sigma). Crushed spleens were washed with ice-cold HBSS+HEPES, centrifuged 500 x g for 5 min before red blood cells were lysed in 1X PharmLyse (BD Biosciences, San Jose, CA USA) for 8 min at room temperature (RT). Lysed cells were washed with HBSS+HEPES, centrifuged as above, resuspended in RPMI-1640 (Sigma), and counted on a Bürker grid hemocytometer.

### Flow Cytometry staining and analysis

Spleen cells (500 000 per mouse), and total cells isolated from dcLNs and brain were each stained separately. Cells were first washed with D-PBS, and centrifuged at 400 x g for 5 min. The supernatant was removed, and then Zombie NIR fixable viability dye (1:1000 BioLegend, San Diego, CA, USA) was added for 15 min at RT. Without washing, CD16/32 FcR block (5 μg/ml, BD Biosciences) was added followed by the appropriate antibody mix. Antibodies used: TCRβ PE-Cy7 (1:100 or 1:200 clone H57-597), CD44 PE (1:300 clone IM7) (both BioLegend); CD8a APC-R700 (1:150 or 1:200, clone 53-6.7), CD69 BV421 (1:100, clone H1.2F3), CD25 BB515 (1:150, clone PC61) (BD Biosciences); CD4 FITC (1:500, clone RM4-5), CD4 eFluor506 (1:500, clone RM4-5), CD8 PerCP eFluor710 (1:300, clone 53-6.7), CD44 APC (1:300 or 1:400, clone IM7), FoxP3 (1:40, clone FJK-16s) (eBioscience Thermo Fisher Scientific, Waltham, MA, USA); CD69 APC (1:20, clone H1.2F3, Miltenyi Biotech). All antibodies were used at titers determined empirically under experimental conditions.

Cells were incubated for 30 min at 4 °C. Afterwards, samples were washed twice in HBSS with 1 % FBS and then run on FACSAriaIII (BD Biosciences) equipped with 488 and 633 nm lasers, or on CytoFLEX S (Beckmann Coulter) equipped with 405, 488, 561 and 638 nm lasers, both with standard configuration. Compensations were made using OneComp and UltraComp Beads for antibody fluorescence (eBioscience Thermo Fisher Scientific) and ArC amine reactive beads for viability dye (Molecular Probes, Eugene, Oregon, USA). Fluorescent-Minus-One (FMO) controls were made to ensure gating. These control samples contained all antibodies except one to display fluorescent spreading error of compensated data in each channel (Roederer, 2002). Data were analyzed using FCSExpress v5 (Denovo Software, Los Angeles, CA, USA) and FlowJo v10.4 (Treestar, Portland, OR, USA). The gating strategy used for the flow cytometry analysis of brain-isolated immune cells is reported in Figure 1.

**Figure 1.**
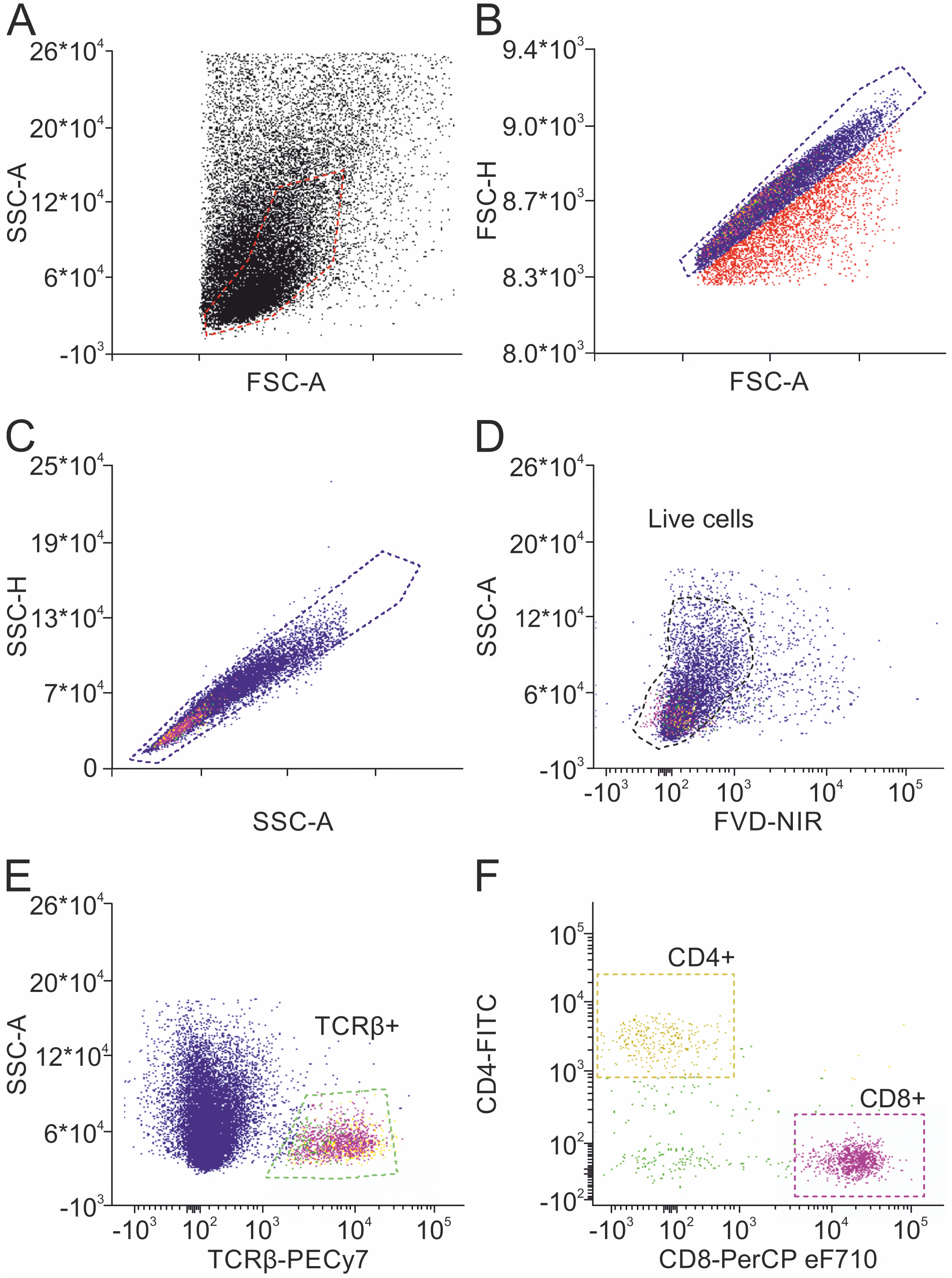
Gating strategy. Flow cytometry analysis scheme showing how isolated immune cells in the brain were gated for live cell analysis. Mononuclear cells were discriminated out from debris (red dotted line) by light scattering properties in a 2D plot showing forward scatter (FSC-A) *vs.* side scatter (SSC-A) (A). These gated cells were analyzed further with Height (-H) and Area (-A) parameters of FSC and SSC to remove cell doublets (B, C). From these single gated events, live cells were defined as negative for near infrared fluorochrome-conjugated fixable viability dye (FVD-NIR) (D), then from these live cells, T cells were identified as positive for TCRβ (E). TCRβ+ lymphocyte subsets were characterized by the expression of CD4 and CD8 cell surface markers (F). Gating was determined using FMOs (see Methods section).

### CD3 immunohistochemical staining

Three mice per genotype were injured and sacrificed 30 days after TBI for the immunohistochemical (IHC) estimation of T lymphocyte localization in the brain. Mice were transcardially perfused with ice-cold NaCl 0.9 % followed by 4 % PFA. Brains were dissected and post-fixed in 4 % PFA by immersion for 24 h at 4 °C. Thereafter, specimens were cryoprotected by incubation in 20 % glycerol (in 0.02 M potassium phosphate-buffered saline (KPBS), pH 7.4) for 48 h, frozen in N-pentane (3 min at −60 °C), and stored at −70 °C until sectioning. Frozen coronal sections were cut 25 μm with a sliding microtome, and collected in solution containing 30 % ethylene glycol, 25 % glycerol in 0.05 M phosphate buffer (PB) and stored at −20 °C until further processing. Three sections per brain (approx. 700 μm apart, encompassing the antero-posterior extension of the lesion,) were used to estimate the localization of CD3+ infiltrating T lymphocytes by IHC. Floating sections were washed in three changes of 1X PBS before being incubated for 1 h at RT in blocking solution (2 % normal goat serum, 1 % bovine serum albumin (BSA) 0.1 % Triton X-100 and 0.05 % Tween20 in PBS). Sections were incubated overnight at 4°C with rat anti-mouse CD3ε in staining buffer PBS with 1 % BSA and 0.05 % Triton X-100 (1:500, clone 17A2, eBioscience Thermo Fisher Scientific). After washing 3x with PBS, sections were incubated with secondary antibody Alexa Fluor 647-or Alexa Fluor 488-conjugated goat anti-rat secondary antibody in above staining buffer for 1 h at RT (1:500, both Thermo Fisher Scientific). Finally, the sections were washed 3x in PBS and 10 min in 1X PB, and mounted on Superfrost Plus slides (Thermo Scientific) with Vectashield (Vector Laboratories, Burlingame, CA, USA). Panoramic photomicrographs of the stained sections were captured using 20X objective with a fluorescence microscope (Zeiss Observer.Z1), and high-resolution Z-stack images were captured using 20X objective with a confocal microscope (Zeiss LSM710). ZEN 2012 software (Carl Zeiss GmbH) was used for image processing.

### MAP2, NeuN and GFAP staining and analysis

Three sections located at bregma level +0,02 mm, −2,06 mm and −4,04 mm (corresponding to the anterior and posterior edges and to the center of the lesion site) were selected from the previously sliced brains and stained for the Microtubule-Associated Protein 2 (MAP2; neuronal dendrites), the neuronal antigen NeuN, and the Glial Fibrillary Acidic Protein (GFAP; Type III intermediate filaments in astrocyte). For immunofluorescence procedure, sections were washed and blocked in blocking solution (4 % BSA, 0,2 % Triton X-100 in PBS) for 1 h at RT, followed by overnight incubation at 4 °C with the following primary antibodies diluted in blocking solution: mouse anti-GFAP (1:500, Sigma G3893), guinea pig anti-NeuN (1:500, Millipore ABN90), rabbit anti-MAP2 (1:300, Abcam ab32454). After washing in PBS, sections were incubated for 2 h at RT with secondary fluorescent antibodies in blocking solution: Alexa Fluor 546-conjugated goat anti mouse (1:250), Alexa Fluor 488-conjugated goat anti rabbit (1:250), Alexa Fluor 633-conjugated goat anti guinea pig (1:200 all from Invitrogen, Thermo Fisher Scientific). Next, sections were washed in PBS before being mounted onto glass slides and coverslipped using Fluoromount-G (Thermo Fisher Scientific).

Image acquisition was performed using Zeiss Axio Observer Z1 microscope, equipped with a Zeiss AxioCam MR R3 camera, mounting a 10x lens to obtain images from whole-brain sections.

Image analysis was performed using ImageJ software. Regions of interest (ROIs) were manually selected on images taken from each stained section. After background subtraction, the mean gray value was measured within each ROI (Clement et al., 2019).

## Statistical analysis

### Data exclusion criteria

We conducted 9 independent experiments, where a total of n = 16 “WT CCI”; n = 12 “WT naïve”; n = 13 “TG CCI” and n = 10 “TG naïve” mice have been analyzed.

Before statistical analysis, brain-derived samples were checked for their quality, based on total T cell recovery. Each sample has been considered independently, and we evaluated the T cell viability and the total number of T cells recovered. Brain samples where T cell viability was below 75 % or the total number of live T cells was below 100 counts were *a priori* excluded from the analyses.

Considering two genotypes (WT and TG) and three experimental conditions (T cells infiltrating the brain tissue ipsilateral to the lesion – “ipsi”; T cells infiltrating the tissue contralateral to the lesion – “contra”, and T cells from naïve brain tissue – “naïve”), a total of n = 12 “WT ipsi”; n= 7 “WT contra”; n = 5 “WT naïve”; and n = 10 “TG ipsi”; n = 7 “TG contra”; n = 9 “TG naïve” were finally used for statistical analyses.

T cell viability > 90 % was used for the quality requirement of spleen and dcLN samples. Moreover, we excluded spleen samples presenting more than 50 % of necrotic tissue (defined as dark red non-perfused area in the spleen). Considering two genotypes (WT and TG) and two experimental conditions (CCI and naïve), a total of n = 13 “WT CCI”; n= 12 “WT naïve”; and n = 11 “TG CCI”, n = 9 “TG naïve” spleens were used for subsequent statistical analyses. Deep cervical lymph nodes have been analyzed in n = 4 “WT CCI” and n = 6 “TG CCI” mice.

### Statistical analysis of brain- and dcLNs-related data

Due to the small amount of T lymphocytes in naïve brains, brain samples were fully acquired on the flow cytometer, and for each population we analyzed both the absolute counts and the percentage referred to the respective parent population. Statistic models were applied considering the nature of our data (counts or percentages) and the experimental groups analyzed. A binomial negative regression was applied to assess statistical differences in the counts of total T cells, of CD4+, and of CD8+ cells between the two genotypes or within the same genotype between independent data. The binomial negative regression considered both genotype and treatment and their interaction. Because data from ipsi and contralateral brain sides are dependent within the same genotype, a linear mixed model was used to evaluate the differences in the total number of CD4+ and CD8+ T lymphocytes between “WT ipsi” vs. “WT contra” or “TG ipsi” vs. “TG contra”. As the data were not normally distributed (Shapiro-Wilk test p-value < 0.05), statistical differences between independent data in CD4+ and CD8+ T cell populations (expressed as percentage of T cells), as well as in the percentages of respective subpopulations expressing CD44 and/or CD69 antigens, were analyzed performing the Kruskal Wallis test. Dependent data within the same genotype (ipsi vs. contra) were analyzed performing the paired samples Wilcoxon signed ranked test. In all tests, Bonferroni correction was used to adjust p-values in multiple comparison.

### Statistical analysis of data from spleen

All data from spleen are expressed as percentage of the parent population. After establishing the normal distribution of the data (as well as skewness and kurtosis by D’Agostino K-squared test), statistical differences were analyzed performing the Kruskal Wallis test or the paired samples Wilcoxon signed ranked test, depending on the nature of the data (independent or dependent), followed by Bonferroni adjustment.

### Statistical analysis of MRI data

The differences in contusion volume and in brain atrophy were analyzed performing the Kruskal Wallis test. Correlation between TBI-related tissue loss and infarct volume was analyzed by Pearson linear regression, after checking for normal distribution of data as described above.

Statistical analyses were performed using R v3.5.3 software/computing environment (The R foundation for statistical computing). All software packages (MASS, psych, agricolae, multcomp and lme4) (Mendiburu, 2019, Revelle, 2018, Bates et al., 2015, Hothorn et al., 2008, Venables and Ripley, 2002) were taken from the Comprehensive R Archive Network mirror sites (CRAN; http://CRAN.R-project.org/package=boot). Significance was accepted at the level of p < 0.05.

## Results

### T cells preferentially infiltrate the cortical areas ipsilateral to the lesion

The presence of infiltrating T lymphocytes in the parenchyma is a signature of brain lesion. At a chronic time point after TBI, we estimated T cell presence in the area of injury and in other brain areas not directly affected by the penetrating injury. For this purpose, we stained brain sections of both WT and TG mice at 30 days post-injury (dpi) for the presence of CD3, a specific marker of T lymphocytes. As expected, T cells are massively present within the boundaries of the injured area (Figure 2A, B; Supplementary Figure 1B). CD3+ cells are also spread throughout the cortical parenchyma, both in proximity to the lesion core (Figure 2C) and in more distal areas ipsilateral to the lesion along the cortical layers. Positive immunostaining was also found along the corpus callosum (Figure 2D; Supplementary Figure 1B) while a minor presence of T cells was observed in the striatum, the hippocampus, and the thalamus ipsilateral to the lesion (Figure 2A). Dim CD3+ signal was present in the contralateral hemisphere, indistinguishable from non-injured mice (data not shown). There was no difference in T cell distribution between WT and TG mice. Unevenly scattered T cells (Figure 2E) and T cell clusters (Figure 2C, D) were both observed within the parenchyma, suggesting the presence of clonal expansion of activated T cells.

**Figure 2.**
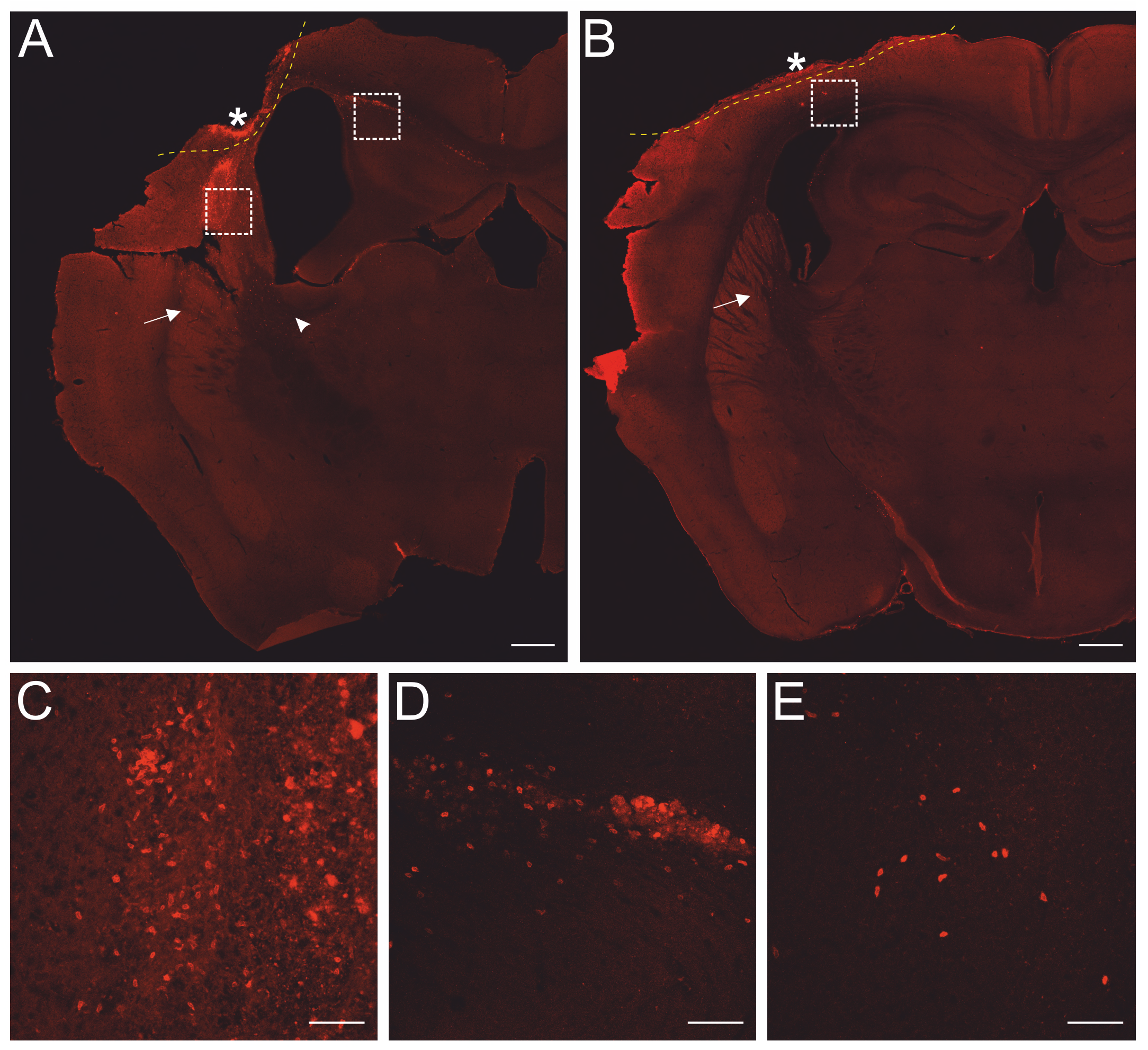
Localization of CD3+ T cells in the perilesional cortices. Representative images of brain sections from WT CCI **(A)** and TG CCI **(B)** mice 30 dpi, stained for anti-CD3ε (T lymphocytes; red). The lesion edges in each section are marked with a segmented yellow line. T cells are present within the lesion (star in A and B), in the perilesional cortex (box in A and panel C) and in the corpus callosum (box in A and panel D). CD3+ cells were also observed in the striatum (arrow in A and B) and in the thalamus (arrow head in A). Both scattered cells and clusters of T cells were found within the parenchyma (C and E, respectively). Panels **(C)** and **(D)** represent a magnification of the areas depicted within the white boxes in A. Panel **(E)** represents a magnification of the area depicted within the white box in B. (A and B, scale bar = 500 μm; C-E, scale bar = 20 μm.)

Next, we decided to quantify and characterize the populations of infiltrating T lymphocytes using flow cytometry, focusing on the neo-cortical areas (cortices and hippocampi), excluding the lesion area, which is characterized by a dysregulated entrance of immune cells (Fee et al., 2003).

Thirty days after brain trauma induction in TG and littermate WT mice, leukocytes were purified separately from the perilesional and the contralateral cortices (or from the cortex of both WT and TG naïve mice). T cells were identified by staining for T cell receptor (TCRβ) and the presence of the co-receptors CD4 and CD8. The acquired count of live T cells in the different experimental conditions is reported in Figure 3. A significant ~10-fold increase of infiltrating T cells was found in both WT (median = 1449; Q3-Q1 = 1692) and TG (median = 1741; Q3-Q1 = 892) mouse brains in the perilesional cortices, compared to corresponding naïve non-injured mice (WT naïve: median = 242; Q3-Q1 = 105; TG naïve: median = 197; Q3-Q1 = 66; for statistical analysis, see Figure 3A). In the cortices contralateral to the lesion, the number of TCRβ+ cells was no different from naïve brains (WT contra: median = 201; Q3-Q1 = 84; TG naïve: median = 239; Q3-Q1 = 155; for statistical analysis, see Figure 3A). No genotype-related differences were observed (Figure 3A).

**Figure 3.**
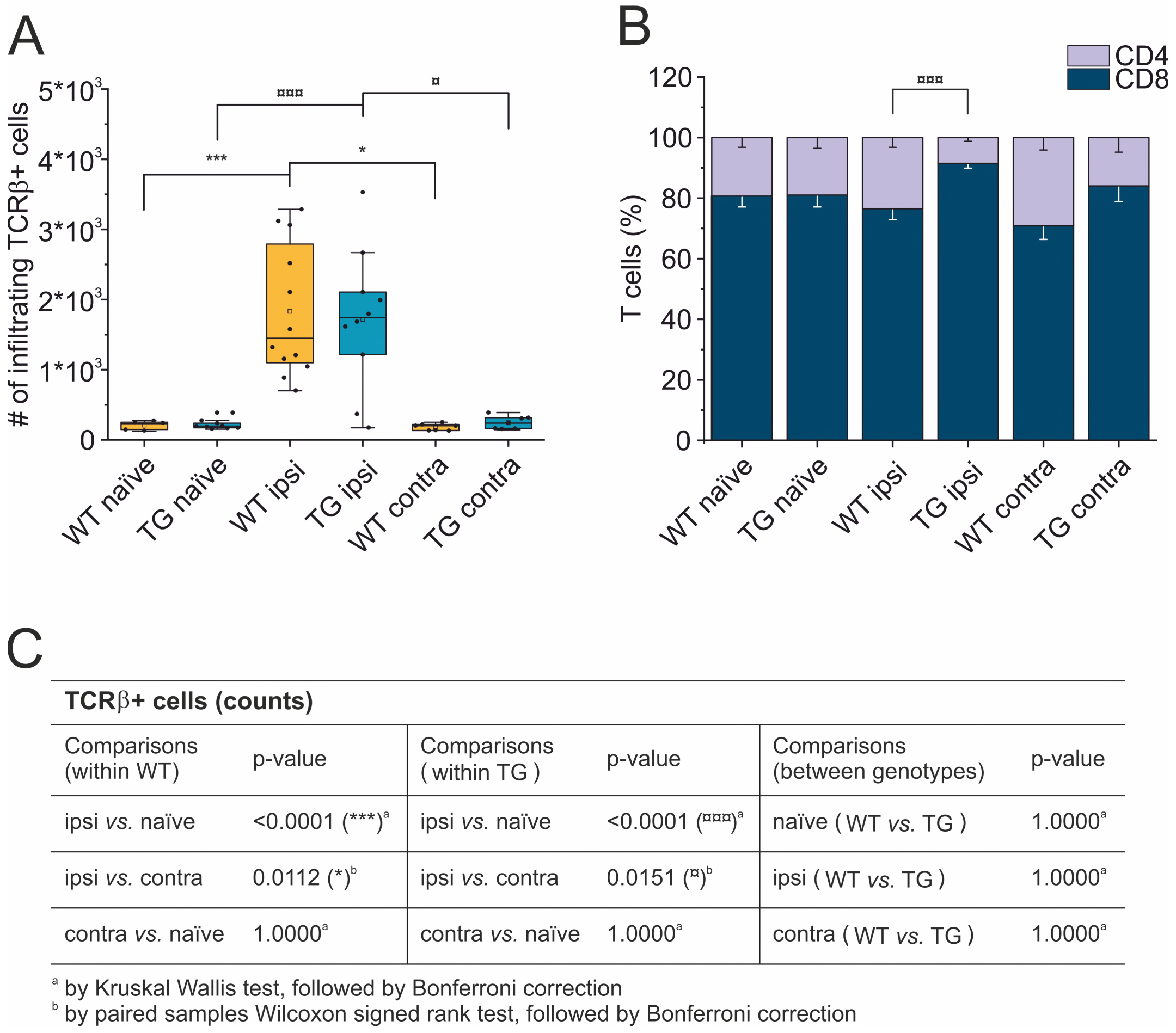
T cell brain infiltration is confined to the perilesional cortices, 30 dpi. Box plot representing the number of infiltrating T cells, defined by expression of TCRβ **(A)** and stacked bargram representing the percentage of CD4+ and CD8+ T cells **(B)** in the brain of WT and TG mice, as analyzed in the perilesional and contralateral cortices (ipsi and contra, respectively), or in intact cortices from respective naïve mice. Independently from the genotype, a significant infiltration of TCRβ+ T cells was observed in the perilesional areas but not in the contralateral hemispheres (comparable to naïve non-injured brains). The majority of brain-infiltrating T cells presented a CD8 phenotype. In the TG CCI mice, there was a significant skew of CD4/CD8 ratio towards CD8+ T cells. Table **(C)** summarizes the results of the statistical analysis in T cell counts between the experimental groups. In (A) boxes represent the 25-75 % value range, including the median value, indicated with the line. Whiskers represent 1.5x standard deviation (SD). □ indicates the mean value. In the stacked bargram, data are presented as mean ± standard error of the mean (s.e.m.). A binomial negative regression or a linear mixed model was applied to assess statistical differences in the counts of total T cells. The Kruskal Wallis test or the paired samples Wilcoxon signed ranked test was used for the analysis of CD4 and CD8 frequency distribution. ¤p < 0.05 and ¤¤¤p < 0.001 vs. TG ipsi. *p < 0.05 and ***p<0.001 vs. WT ipsi. In all tests, Bonferroni correction was used to adjust p-values in multiple comparisons.

### Perilesional-infiltrating T cells have a predominant CD8+ phenotype, and the lack of a functional lymphatic system depresses the T cell CD4-mediated response

We next analyzed the CD4:CD8 ratio within the infiltrating T cells (Figure 3B) and found a prevalence of CD8+ T cells in all the experimental conditions, regardless of the presence of brain injury. However, limited to the perilesional cortex of TG mice, we detected a significant skew of the CD4:CD8 ratio towards CD8+ cells (CD4:CD8 ratio TG ipsi = 0.097±0.053; WT ipsi = 0.350±0.197; ChiSq: 8.836, mean ranks: 5.50/13.27, p = 8e-04), while the ratio in the contralateral cortex did not differ between the two genotypes (CD4:CD8 ratio TG contra = 0.221±0.247; WT contra = 0.456±0.212; ChiSq: 2.469, mean ranks: 5.43/8.83, p = 0.120). To better understand how the lack of mLVs affects the T cell-mediated neuro-immune response, we analyzed both the absolute numbers of CD4 and CD8 subpopulations and their relative frequency. Data analysis shows a reduction of the total number of CD4+ T cells infiltrating the perilesional cortices of TG (median = 106; Q3-Q1 = 156) compared to WT mice (median = 245; Q3-Q1 = 218; ex. coef.: −0.82, p = 0.033 TG ipsi vs. WT ipsi) (Figure 4A). No differences were observed in the absolute number of infiltrating CD8+ T cells between the genotypes (Figure 4B). Despite no differences in absolute numbers of both CD4 and CD8 populations in the contralateral cortices of injured WT and TG mice, we found a significant reduction in the frequency of CD4+ T cells in transgenic mice (TG contra = 12.04±8.47 %; WT contra = 23.59±9.52 % of T cells; ChiSq: 3.931, mean ranks: 5.29/9.71, p = 0.042) and a relative frequency increase of CD8+ T cells (Figure 4C, D). These data are in line with previous studies indicating that the CD4-mediated neuro-immune response is mainly induced within the dcLNs (Thomas, D. L. et al., 2008, Harling-Berg et al., 1999). As mLVs are involved in the drainage of solutes from the interstitial and cerebro-spinal fluids mainly to the dcLNs (Aspelund et al., 2015, Louveau et al., 2015), it is possible to conceive that their absence in TG mice can affect the priming of the evoked neuro-immune response, resulting in a specific impairment of CD4+ T cell activation. The analysis of the T cell subpopulations in the dcLNs indeed revealed a marked difference between the two genotypes, supporting the role of mLVs in the definition of the neuro-immune response. We found a significantly lower number of T cells in the dcLNs of the TG-CCI mice (median = 73542; Q3-Q1 = 21342) compared to their WT-CCI littermates (median = 220434; Q3-Q1 = 88745; p = 0.006), which had a higher frequency of CD4+ T cells (TG CCI = 63.98±5.67 %; WT CCI = 51.40±1.93 % of T cells; ChiSq: 6.545, mean ranks: 7.50/2.50, p = 0.0017) (Supplementary Figure 2). Within the CD4+ T cell subpopulation in the TG mice, cells have predominantly a CD44^hi^CD69^neg^ phenotype, while in the WT mice the predominant population is CD44^int^CD69^neg^ (Supplementary Figure 2). No differences were found in the frequency of Tregs.

**Figure 4.**
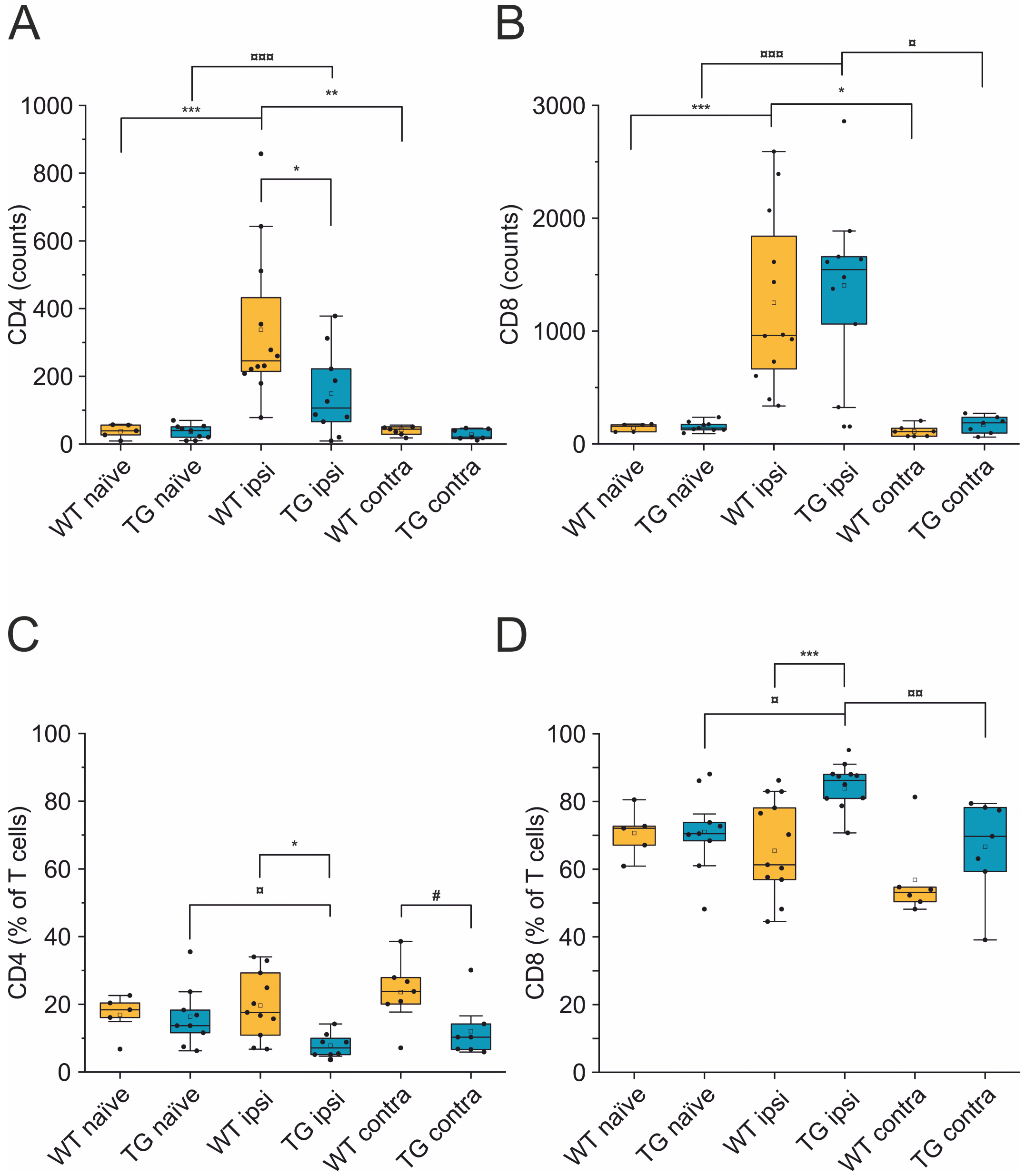
The number of CD4+ but not of CD8+ T cells is reduced in the brain of K14-VEGFR3-Ig mice after TBI. Box plots representing the number and frequency of CD4+ T cells (**A** and **C**, respectively) and CD8+ T cells (**B** and **D**, respectively), in the brain of WT and TG mice, as analyzed in the perilesional and contralateral cortices (ipsi and contra, respectively), or in intact cortices from naïve mice. A drastic reduction in the number of CD4+ T cells was found in TG mice after injury. A binomial negative regression or a linear mixed model was applied to assess statistical differences in the counts of CD4+ and CD8+ T cells. The Kruskal Wallis test or the paired samples Wilcoxon signed ranked test was used for the analysis of frequency distribution. *p < 0.05; **p < 0.01 and ***p < 0.001 vs. WT ipsi. ¤p < 0.05; ¤¤p < 0.01 and ¤¤¤p < 0.001 vs. TG ipsi. #p < 0.05 vs. WT contra. In all tests, Bonferroni correction was used to adjust p-values in multiple comparisons. For box plot explanation, refer to the legend of Figure 3.

The presence of CD8+ T cells in the perilesional cortices (together with the presence of T cell clusters, as shown by IHC staining) suggests a cytotoxic role for the infiltrating T cells at this time point. However, different subpopulations of CD8+ and CD4+ T cells exist, with specific and opposing functions. In addition, we characterized both the CD8+ and CD4+ subpopulations for the surface expression of the antigens CD44 (a memory and activation marker) (Ponta et al., 2003, Budd et al., 1987) and CD69 (an activation and tissue retention marker) (Ziegler et al., 1994). In the perilesional cortex of both WT and TG mice, CD8+ T cells had a predominant CD44^hi^CD69+ phenotype (69.78±22.85 % and 72.05±19.95 % of CD8+ T cells, in WT ipsi and TG ipsi, respectively) (Figure 5A, B and Supplementary Table 2). In the mouse, the expression of CD69 together with high levels of CD44 define a specific subpopulation of T cells called mature resident memory T cells (T_RM_), which are generated and persist in the tissue at the site of a primary infection (Topham and Reilly, 2018, Gebhardt et al., 2009) and provide a first and powerful line of adaptive cellular defense.

**Figure 5.**
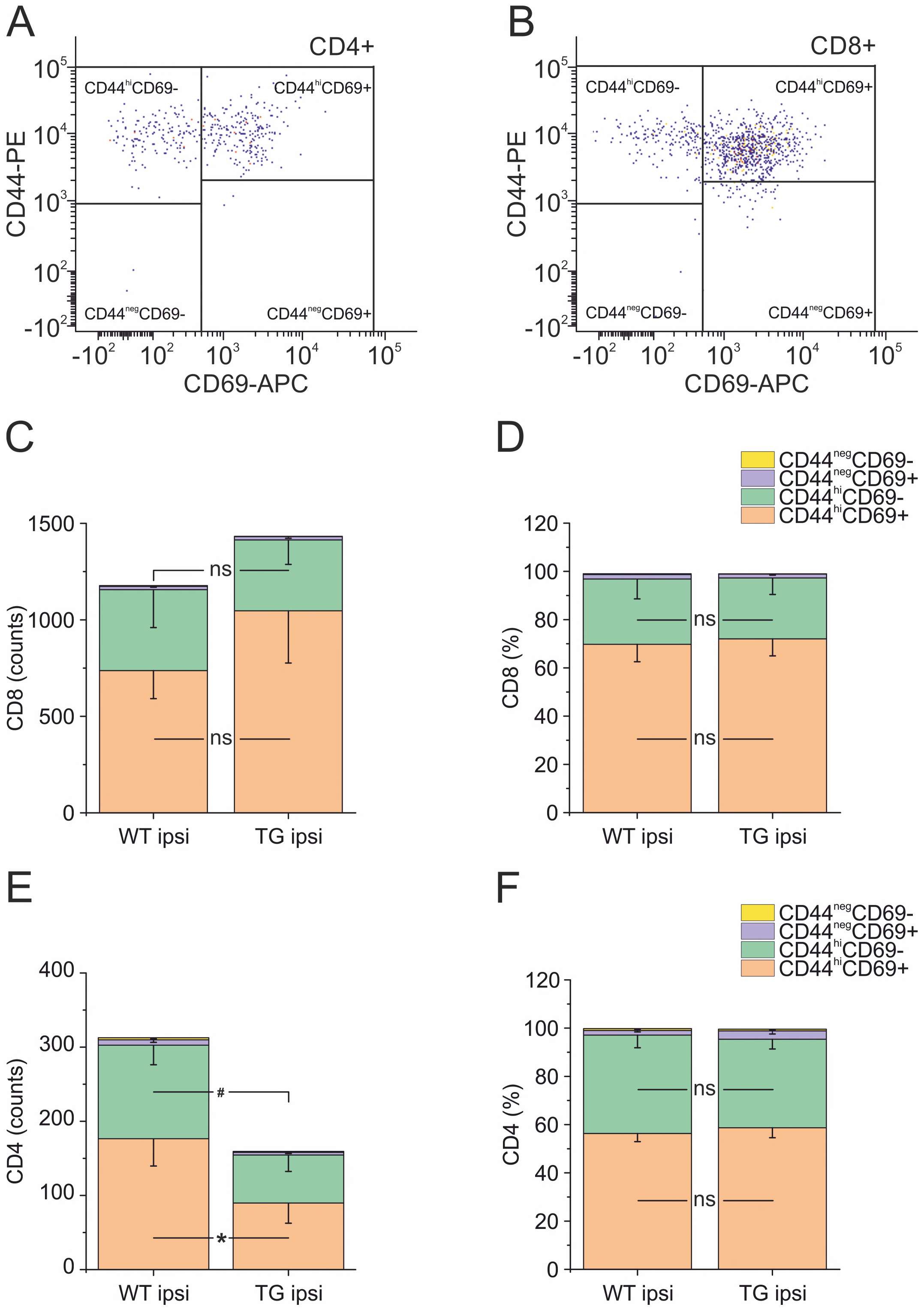
Analysis of CD69 and CD44 T cell activation and memory markers in CD4+ and CD8+ subpopulations. Pseudocolor dot plots **(A)** and **(B)** represent gated subpopulations CD69 vs. CD44 of CD4+ and CD8+, respectively. Stacked bargrams in **(C)** and **(D)** show respectively the counts and frequencies of CD8+ T cell subpopulations, as analyzed in the perilesional cortices of WT and TG mice. No significant differences in CD8+ subpopulations were found between genotypes. In CD4+ subpopulation, instead, we observed a significant reduction in the counts of CD44^hi^CD69+ and CD44^hi^CD69-subpopulations **(E)**, in K14-VEGFR3-Ig compared to WT mice. However, no differences were observed in the different subpopulation frequencies **(F)**. Data are presented as mean ± s.e.m. A binomial negative regression was applied to assess statistical differences in the counts of total T cells between WT ipsi and TG ipsi. The Kruskal Wallis test was used for the analysis of frequency distribution. #p < 0.05; *p < 0.05 vs. WT ipsi.

The second-highest expression of a CD8+ subpopulation (representing 27.07±26.10 % in WT and 25.24±18.85 % in TG mice) had a CD44^hi^CD69-phenotype, characteristic of effector memory T cells (Topham and Reilly, 2018). The presence of other CD8+ subpopulations among perilesional infiltrating T cells was negligible. No genotype-related difference was found.

Among CD4+ perilesional infiltrating T cells, we found a similar frequency of CD44 and CD69 expressions, with a slight prevalence of CD44^hi^CD69+ over CD44^hi^CD69-T lymphocytes (Figure 5C, D and Supplementary Table 2) in both genotypes. The overall frequency distribution of the different subpopulations was identical between the two genotypes.

### Cortical lesion is similar in K14-VEGFR3-Ig mice and in their WT littermates

Analyses of MRI images acquired 21 days after TBI induction revealed a T2 intensity increase in the ipsilateral hemisphere. The increase of T2 intensity was observed in parietal-temporal cortices, mainly involving the somatosensory and visual cortices (Figure 6A), expanding in a few cases to the underlying hippocampus. No significant change of T2 intensity was found between the two genotypes. In the WT CCI group the contusion volume was 4.53±1.33 %, and 4.09±2.00 % in the TG CCI animals (ChiSq: 0.579, mean ranks: 8.71/10.75, p = 0.463) (Figure 6C). Relative brain atrophy was 2.42±1.09 % in WT CCI mice and 2.00±1.26 % in TG CCI mice (ChiSq: 1.400, mean ranks: 8.00/11.17, p = 0.248) (Figure 6D). Correlation between contusion volume and relative brain swelling was compared in transformed data analyzed by linear regression. When considering the individual values independent of the genotype, the contusion volume values significantly correlated with the values of relative brain atrophy (r = 0.57; p = 0.023) (Figure 6E). No significant correlation was found between the contusion volume and the mean value of the brain atrophy in both the TG CCI group (r = 0.74; p = 0.064), and in the WT CCI mice (r = 0.37; p = 0.331).

**Figure 6.**
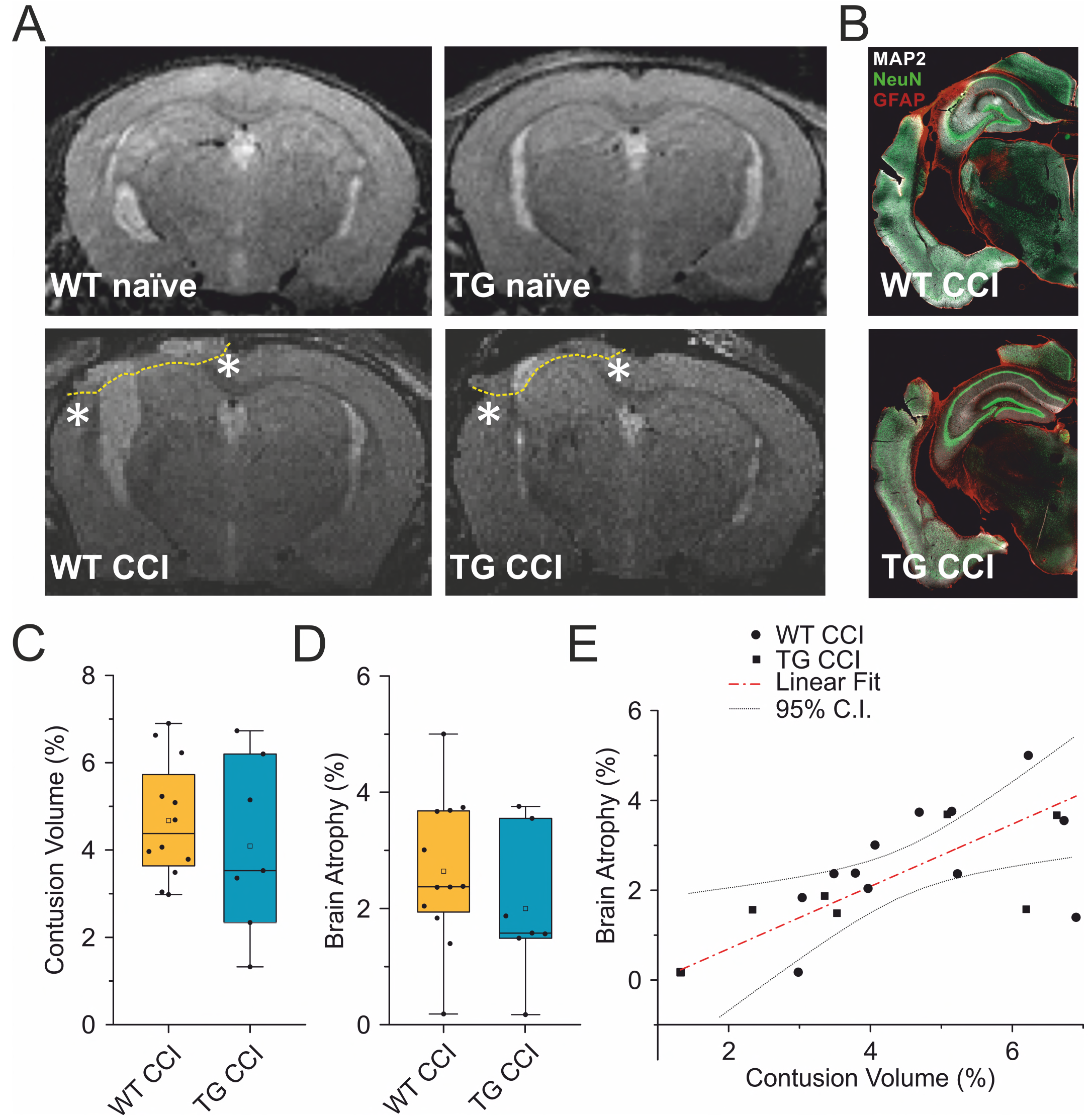
TBI-induced lesions does not differ between the two genotypes, as inferred by the analysis of MRI at 21 dpi. **(A)** Representative MR images of WT naïve, WT CCI, TG naïve and TG CCI brains. Perilesional cortices in WT CCI and TG CCI brains are marked with stars. **(B)** Representative images of WT CCI and TG CCI brains stained for MAP2, NeuN and GFAP at 30 dpi. No differences in neuronal damage or in neuroinflammation were visible between the two genotypes. Box plots in **(C)** and **(D)** illustrate the genotype effect on the percentage of contusion volume and of brain atrophy, respectively, over the volume of the hemisphere ipsilateral to the lesion. No significant differences were observed between K14-VEGFR3-Ig and WT mice. For the definitions of the contusion volume and of brain atrophy see the main text. **(D)** When considering the contusion volume and the brain atrophy independently from the genotype, we found a direct correlation between the two parameters. The Kruskal Wallis test was used for the analysis of infarct volume and of tissue loss between the two genotypes. CI: 95 % confidence interval. For box plot explanation, refer to the legend of Figure 3.

It must be noted that we have identified the lesion size as the hyper-intense signal in the cortical area observed in the T2 weighted images. Our analysis, albeit clinically relevant, suffers from a lack of spatial definition and is affected mostly by the formation of the cyst at the site of injury (Maegele et al., 2015, Immonen, R. et al., 2010). Therefore, subtle although significant differences in the lesion size can be underestimated. However, the analysis of MAP-2 staining in the brain of the WT CCI and TG CCI animals, used for the evaluation of T cell presence in the injury area, confirmed the MRI results and did not show any genotype-related differences (Figure 6B).

### K14-VEGFR3-Ig mice present a peripheral lymphopenia, which is exacerbated after TBI

Alterations of systemic immunity are frequent in TBI patients. We analyzed the levels and the frequency of different T cell subpopulations in the spleen of WT and K14-VEGFR3-Ig mice, one month after TBI induction. As previously described (Thomas, S. N. et al., 2012), K14-VEGFR3-Ig mice show a moderate lymphopenia compared to littermate WT mice (percentage of T cells over live cells in WT naïve: 37.26±7.67 %; vs. TG naïve: 19.69±4.96 %; ChiSq: 14.746, mean ranks: 5.00/15.50, p = 1e-04) (Figure 7A). Contrary to what was observed in the brain, the systemic lymphopenia in the K14-VEGFR3-Ig genotype corresponds to a relative frequency reduction in peripheral CD8+ T cells (TG naïve = 25.75±3.61 %; WT naïve = 42.70±4.17 % of T cells; ChiSq: 14.727, mean ranks: 5.00/15.50, p = 1e-04) (Figure 7B). In TG mice, but not in WT mice, we found a significant reduction in the total T cell frequency after TBI (WT CCI: 33.68±6.99 %; TG CCI: 14.23±2.87 % of live cells; ChiSq: 7.695, mean ranks: 7.18/14.55, p = 0.003 TG CCI vs. TG naïve) (Figure 7A), confirming that TG mice present an impaired immune response, which relates to the alterations in the lymphatic system. Analysis of the activation markers show a different expression in both CD4+ and CD8+ subpopulations between WT and TG mice, which is trauma independent. Both TG naïve and TG CCI mice, indeed, showed an increased frequency of memory T cells (CD4+CD44^hi^CD69+, CD4+CD44^hi^CD69- and CD8+CD44^hi^CD69+, CD8+CD44^hi^CD69-; for statistical analysis, see Supplementary Table 3) (Figure 7C, D).

**Figure 7.**
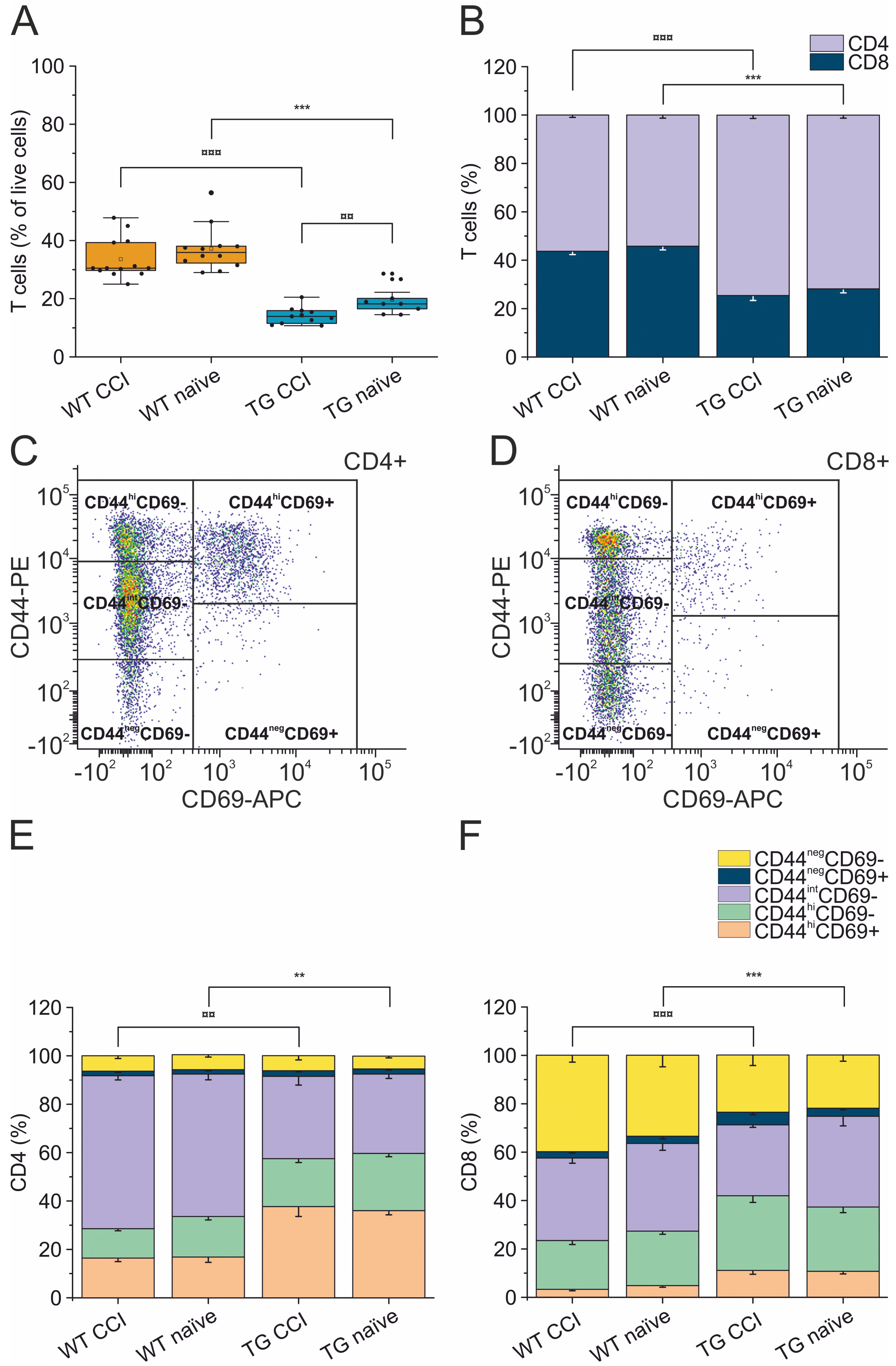
Peripheral immune response in the spleen. The percentages of T cells in the spleen of WT naïve and CCI mice and of TG naïve and CCI mice are presented in the box plot in panel **(A)**. Stacked bargrams in **(B)** represent the relative percentages of CD4 and CD8 in T cell population, in WT and K14-VEGFR3-Ig mice. K14-VEGFR3-Ig mice present a drastic reduction of T cells compared to WT littermates, due to a decrease in CD8+ T cell frequency. **(C, D)** Representative pseudocolor dot plots and gating strategies for CD4+ and CD8+ T cell subpopulation analysis, respectively. Stacked bargrams in **(E)** and **(F)** show respectively the frequencies of CD4+ and CD8+ T cell subpopulations, as analyzed in WT and TG mice. Significant differences in the frequencies of both CD4+ and CD8+ subpopulations have been observed. The Kruskal Wallis test or the paired samples Wilcoxon signed ranked test was used for the analysis of frequency distribution. ¤¤p < 0.01 and ¤¤¤p < 0.001 vs. TG CCI. **p < 0.01 and ***p < 0.001 vs. WT naïve. In all tests, Bonferroni correction was used to adjust p-values in multiple comparison. For box plot and stacked bargram explanation, refer to the legend of Figure 3.

## Discussion

The results of this study show the effects of the deficiency of a functional CNS lymphatic system on the expansion of brain-resident memory T cells as a result of a single, moderate TBI.

Mounting evidence implicate a sustained modulation of T lymphocyte-mediated immune response following TBI, both in patients (Dressler et al., 2007, Hausmann et al., 1999, Holmin et al., 1998) and in animal models of brain injuries (Bai et al., 2017, Braun et al., 2017, Erturk et al., 2016, Kelso and Gendelman, 2014, Jin et al., 2012).

A recent publication from Daglas and colleagues characterized for the first time the T cell-mediated immune response in a chronic animal model of TBI, highlighting the role of cytotoxic CD8+ T cells in the progression of TBI pathology (Daglas et al., 2019).

Our data confirm the previous findings, showing a sustained accumulation of CD8+ T lymphocytes, restricted to the non-damaged cortical areas surrounding the lesion and to the underlying corpus callosum, already at 30 dpi *(i.e.,* the early chronic phases after TBI). Moreover, we expand the current knowledge characterizing the phenotype of the accumulating lymphocytes as resident memory T cells. This suggests a direct in-situ activation of the T cell-mediated immune response. We speculate that this chronic activation is responsible for the progression of TBI pathology.

We also found that the congenital lack of the meningeal lymphatic system affects the polarization of the TBI-elicited neuro-immune response, mainly resulting in the downregulation of CD4+ T cell subpopulation. We finally found that the adaptive neuro-immune response is prompted even in the absence of a systemic immune reaction.

Specifically, our findings suggest that at early chronic time points after TBI: 1) immune response in the brain is principally mediated by putative T_RM_ CD8+ cells; 2) the CNS lymphatic system is essential to modulate the specific neuro-immune response; 3) the response of peripheral T lymphocytes does not correlate with the neuro-immunological state of the brain.

Brain trauma results in two phases of tissue injury. The primary injury which is a direct result of the mechanical impact to the brain, is characterized by the activation of the innate immune response and the release of excitotoxic agents. During this acute phase, a massive and dysregulated brain-infiltration of T cells has been reported (Czigner et al., 2007, Clausen et al., 2007). This infiltration is presumably confined to the area of the lesion, since we observed a limited number of infiltrating T cells in the perilesional non-injured areas, three days after TBI induction (Supplementary Figure 2). A secondary tissue damage, resulting in a diffuse and long-lasting injury, usually develops after months/years from the primary injury (Yasmin et al., 2019, Graham and Sharp, 2019, Immonen, R. J., Kharatishvili, Grohn et al., 2009). This is characterized by additional neurodegeneration developing independently from the mechanical trauma and by the formation of a fibrotic scar tissue in the injured area (Fernandez-Klett and Priller, 2014) (Figure 6B). It has been recently suggested that the development of secondary injuries is sustained by activated memory CD8+ T cells (Daglas et al., 2019). In a CCI mouse model, the authors observed that the modulation of the cytotoxic lymphocytes resulted in the reduction of the lesion size and in the improvement of the neurological outcomes analyzed 32 weeks after injury.

In similar experimental conditions, we observed that CD8+ T lymphocytes with a CD44^hi^CD69+ phenotype are already present in the perilesional areas (but not in the correspondent contralateral cortices) one month after TBI. Since CD69 is an early marker of T cell activation (Ziegler et al., 1994) and inhibits tissue egression (Gebhardt et al., 2009), our data suggest a localized activation of the resident memory CD8+ subpopulation restricted to the areas surrounding the primary lesion. In the case of TBI, CD44^hi^CD69+ T_RM_ cells may represent the population designated to defend the non-injured brain from possible infective agents penetrating through the lesion. However, within the chronic neuro-inflammatory environment observed in the perilesional areas (Figure 6B), T_RM_ can expand and activate in a dysregulated way. This contributes to the cytotoxic immune response, which characterizes the chronic phases of TBI pathology. This hypothesis is supported by the data reported by Daglas and colleagues (Daglas et al., 2019), indicating that the perilesional infiltrating CD8+ T cells express and release effector cytokines (Granzyme B and IFNγ). Further studies are required to determine if this adaptive response is antigen specific, and if secondary lesions are the result of an autoimmune-like sequelae of events.

Neuro-immune responses are mainly elicited in the deep and superficial cervical lymph nodes (Cserr et al., 1992, Harling-Berg et al., 1999, Thomas, D. L. et al., 2008, de Vos et al., 2002, Urra et al., 2014), which are the main receivers of the mLVs. Therefore, the meningeal lymphatics represent an integrated component in the neuro-immune response (Louveau et al., 2015), and their functional impairment can affect its priming following TBI.

We addressed this hypothesis by inducing TBI in a transgenic mouse model of congenital lymphedema. K14-VEGFR3-Ig mice, expressing soluble VEGFR-3-Ig (Makinen et al., 2001), present alterations in the development of the lymphatic system, resulting in defective growth of mLVs and in sclerotic dcLNs (Antila et al., 2017, Aspelund et al., 2015). This phenotype has been confirmed in our experimental animals.

We found that the neuro-immune response in the K14-VEGFR3-Ig mice significantly differs from the response observed in WT mice after TBI, suggesting that the functional defect in the CNS lymphatic system directly affects the CNS regional immune regulation and modulates the transition between the initial and secondary immune response after TBI. This hypothesis is supported by the observation that the initial T cell infiltration in the perilesional areas (as determined at 3 dpi) is similar in the two genotypes (Supplementary Figure 2), while at 30 dpi there is a marked decrease in the CD4+ T cell frequency specifically in TG mice. This results in the polarization of the neuro-immune response towards CD8+ cytotoxicity, possibly aggravating TBI outcomes as recently suggested (Daglas et al., 2019).

Interestingly, in chronic TBI animals, the analysis of the T cell subpopulation in the CNS-draining dcLNs also showed a marked difference between the two genotypes. CD4+CD44^hi^CD69^neg^ T cells were the predominant subpopulation in TG mice, and CD4+CD44^int^CD69^neg^ T cells were predominant in WT mice (Supplementary Figure 2). It has been suggested that CD4+CD44^int^ T cells could represent the fraction of central memory T helper cells expressing IFN-γ, while CD4+CD44^hi^ would preferably be effector memory cells with a Th17 phenotype (Schumann et al., 2015, Gasper et al., 2014). A Th1/Th17 response has a role in CNS autoimmune diseases (Kebir et al., 2007) and can enhance the cytotoxicity of CD8+ T cells (Daglas et al., 2019, Braun et al., 2017). This would partially explain the direct correlation we found between the frequency of CD4+ T cells and the brain atrophy in TG mice but not in WT littermates (Supplementary Figure 3). However, the panel of antibodies we used for T cell characterization does not allow us to distinguish between the different CD4+ T helper populations *(i.e.,* Th1, Th2 or Th17) without speculation.

Our data suggest that the functional impairment of mLVs observed in K14-VEGFR3-Ig mice modulates the activation of the adaptive neuro-immune response in the downstream dcLNs. However, we cannot exclude other mechanisms in K14-VEGFR3-Ig mice that could modulate the neuro-immune response. For instance, lymphatic vessels play a direct role in the maturation of T cells, and dysfunction of the lymphatics leads to the persistence of immune cells and mediators in tissues, resulting in a chronic inflammation and tissue damage (Tsunoda, 2017). It is conceivable, therefore, that the congenital lack of mLVs in the K14-VEGFR3-Ig mice can affect both the type of the elicited neuro-immune response and its resolution.

Our hypothesis that the chronic cytotoxic response is mediated by T_RM_ cells, and not by circulating T lymphocytes which infiltrate the brain, has important clinical implications. TBI patients generally present a delayed secondary immunodeficiency (CNS injury-induced immunodepression, CIDS) (Meisel et al., 2005, Mazzeo et al., 2006), which is accompanied by an increased susceptibility to systemic infections and is associated with declining neurological outcome and increased mortality.

Analysis of our data suggest that neuro-immune reaction can be elicited in the CNS even in the presence of a systemic congenital lymphopenia (as observed in K14-VEGFR3-Ig mice), excluding a correlation between the extent of brain infiltration and the level of T cells in the periphery (Supplementary Figure 3). This observation has potential clinical implications, because patients with CIDS could at the same time present a sustained adaptive immune response localized in the brain. Immunomodulatory therapies directly targeting the brain-resident memory T cells could benefit TBI patients without affecting their already compromised systemic immune system.

Therapeutic approaches aimed at downregulating the adaptive immune response after TBI have been tested before (Weckbach et al., 2012) with no improvement on the neurological outcome, leading to the hypothesis that the adaptive immune response after brain injuries can have a beneficial activity (Schwartz and Raposo, 2014, Moalem et al., 1999). However, it is important to note that these studies focused on the manipulation of the early wave of T cell infiltration after TBI. Our findings, together with recently published data, indicate that the chronic immune response is the target for the development of specific therapies for the treatment of TBI patients. This includes modulating the progression of the secondary injuries and opening the way to new studies in this direction.

### Limitation of the study

We are aware that this study presents several limitations and further studies are needed to both understand the role of CD8+ T cells in TBI pathology, and the role of mLVs in the modulation of the neuro-immune response. A major limitation stems from the use of TG mice with a congenital and global deficiency in the mLVs. This results in a compromised peripheral immune response, as previously demonstrated (Thomas, S. N. et al., 2012) and confirmed by our spleen data. In their paper, however, Thomas and colleagues reported a delayed but robust CD8-mediated response to peripheral immunization and impaired tolerance. In a similar fashion, we have found an increase in the CD8+ T cell response to putative brain-derived antigens. These data confirm the pivotal role of lymphatic vessels in the modulation of the adaptive immune response and support the hypothesis that the elicited cytotoxic response can escape the intrinsic brain tolerance. Nevertheless, this hypothesis needs to be confirmed in different models that would study the effects of local partial deletions of the mLVs on the activation of the neuro-immune response.

Another limitation of our study is the lack of difference in lesion size between K14-VEGFR3-Ig mice and their WT littermates despite the increase in the number of cytotoxic T cells. As discussed previously, this could be due to limitations in our analytical approach. However, it is also possible that although triggered by cytotoxic T cells, secondary neurodegeneration and associated behavioral correlates may appear at a later time point than the one analyzed in this study. Specific analyses should be conducted in the K14-VEGFR3-Ig mice to assess the long-term effects of mLV deficits on the progression of TBI pathology.

## Conclusions

Our study investigated the phenotype of T lymphocytes infiltrating and persisting in the brain after TBI, pointing to the activation of the CD8+ resident memory T cells in the early chronic response.

Our findings also support the importance of mLVs and dcLNs in maintaining brain immuno tolerance. We, therefore, propose that the modulation of the neuro-immune response via the CNS-lymphatic system, or by directly targeting the brain-resident memory T cells, could offer therapeutic strategies for the treatment of TBI patients.

## Acknowledgments

This study has been supported by the Academy of Finland (Academy of Finland research Fellowship #309479/2017 – FMN; Academy of Finland research Project #314498 – KA), by the Jane and Aatos Erkko Foundation (JK and KA), by the Finnish Brain Foundation Terva Program (KA) and by European Research Council (ERC) under the European Union’s Horizon 2020 research and innovation programme under grant agreement #743155 (KA).

For their help with MRI sequences, the authors would like to thank Mikko Kettunen and Riikka Immonen from Biomedical Imaging Unit, National Bio-NMR facility, A.I.Virtanen Institute for Molecular Sciences, University of Eastern Finland. Authors thank also Carlton Wong, Flavia Scoyni and Bengisan Dvirick for their contribution in performing experiments. Finally, authors thank Dr. Nicola Marchi for his contribution in discussing the conclusions of this work.

## Author Contributions

**SW**: Methodology, Investigation, Validation, Data Curation, Writing – Review and Editing; **MV**: Investigation, Data curation, Formal analysis; **BG**: Software, Formal analysis; **AV**: Investigation, Formal analysis; **MHK**: Investigation, Writing – Review and Editing; **SA**: Resources, Writing – Review and Editing; **KA**: Supervision, Funding acquisition; **JEK**: Supervision, Funding acquisition; **FMN**: Conceptualization, Methodology, Validation, Writing, Supervision, Funding acquisition.

## Data Availability

The raw data supporting the conclusions of this manuscript will be made available by the corresponding author, upon reasonable request, to any qualified researcher.

