## Supplementary for "The CNS lymphatic system modulates the adaptive neuro-immune response in the perilesional cortex in a mouse model of traumatic brain injury"

### Supplementary Figure 1

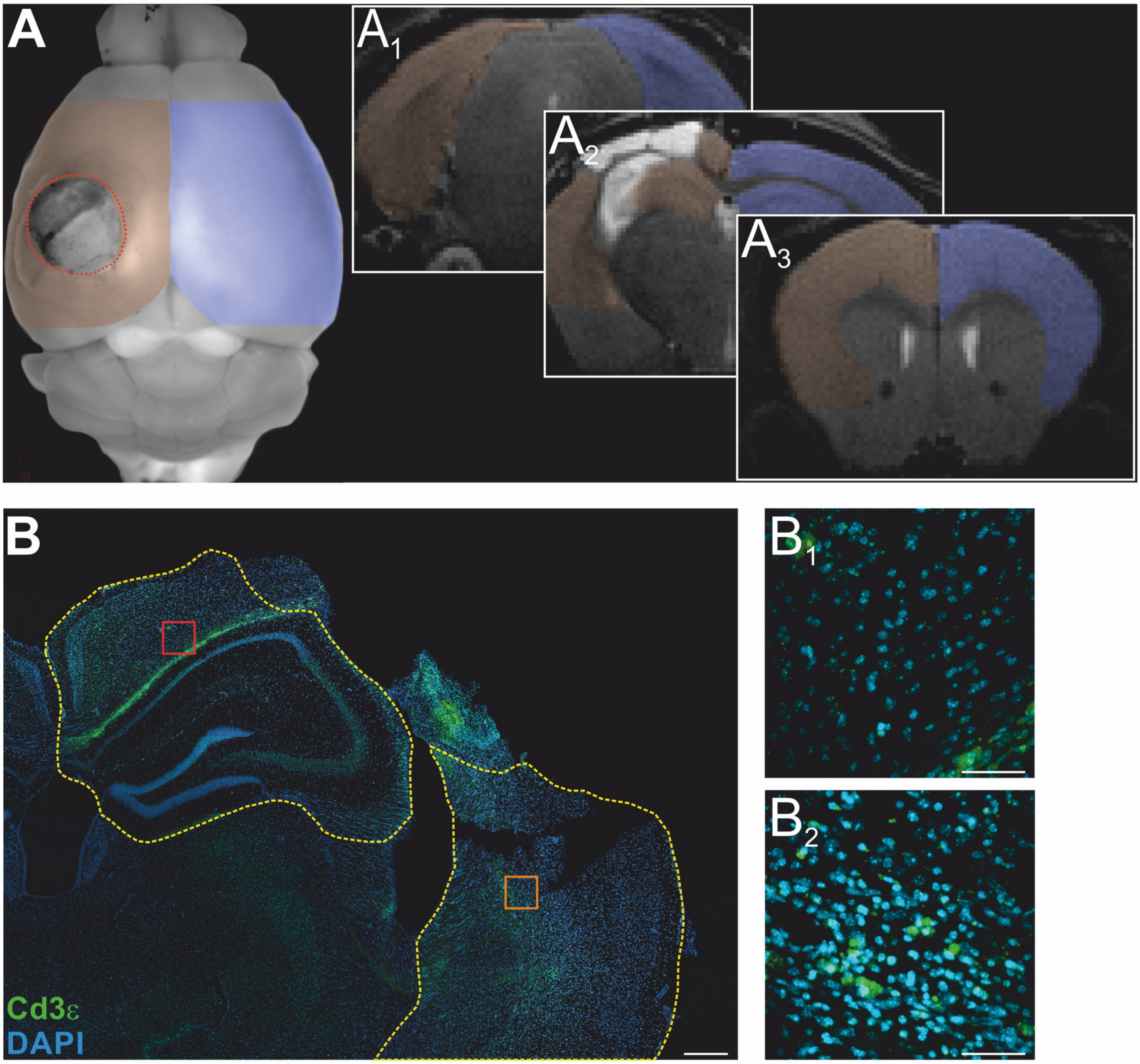


**Supplementary Figure 1:** Representative macrophotograph **(A)** and MR images **($A_{1\text{-}3}$)** of injured brain. Colored ROIs correspond to the perilesional (orange) and contralateral (blue) areas analyzed in the study. Panels $A_1$ and $A_3$ represent respectively the first caudal and the last rostral levels included in the study. Lesion area (delimited by the red dotted line in panel A) is clearly identifiable: to avoid peripheral T lymphocyte contamination, tissue from the lesion area was carefully excised and not used for further analyses. **(B)** CD3+ T cell staining in the cortical areas adjacent to the lesion (perilesional cortex). Areas delimited by the yellow dotted line have been carefully excised and resident T cells isolated. Panels $B_1$ and $B_2$ are the 40X confocal magnification of the areas included in the red and orange boxes in B (respectively). Scale bars: 500 µm (B) and 50 µm ($B_1$ and $B_2$)

Supplementary Figure 2

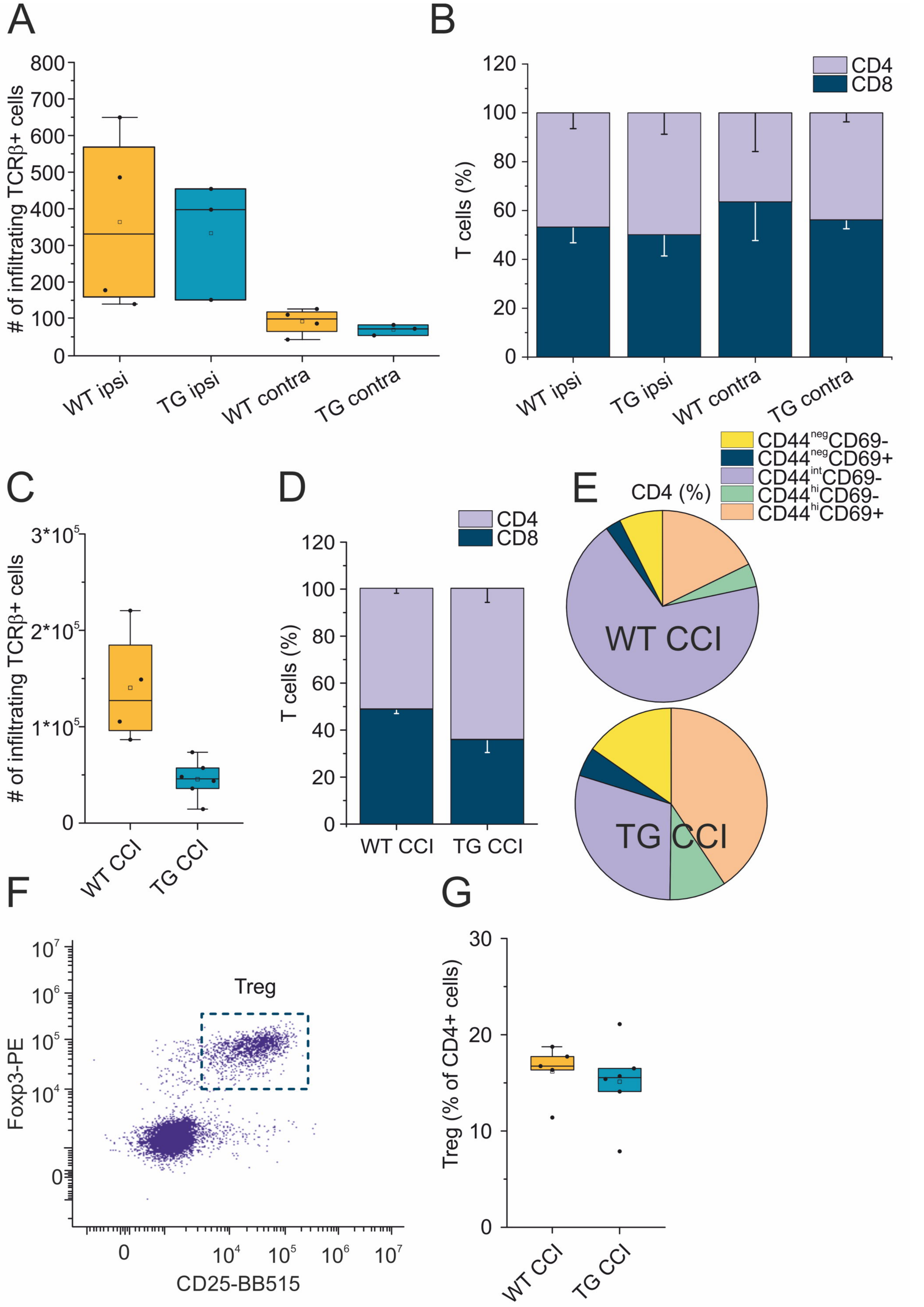


**Supplementary Figure 2: (A, B)** ***T cell infiltration in the brain, 3 dpi*** - Box plot representing the number of infiltrating T cells (A) and stacked bargram representing the percentage of CD4+ and CD8+ T cells (B) in the brain of WT CCI and TG CCI mice. **(C-G)** ***Analysis of T cell populations in the dcLNs, 30 dpi -*** Box plot in (C) and stacked bargram in (D) represent respectively the total number and the CD4:CD8 ratio of T cells in the dcLNs (left and right side combined for each mouse). Pie charts in (E) represent the frequencies of CD4+ T cell subpopulations. (F) Representative dot plot and gating strategy for the analysis of the CD4+FoxP3+CD25+ Treg cell subpopulation. (G) Box plot representing the percentages of Tregs in the dcLNs of WT CCI and TG CCI mice.

### Supplementary Figure 3

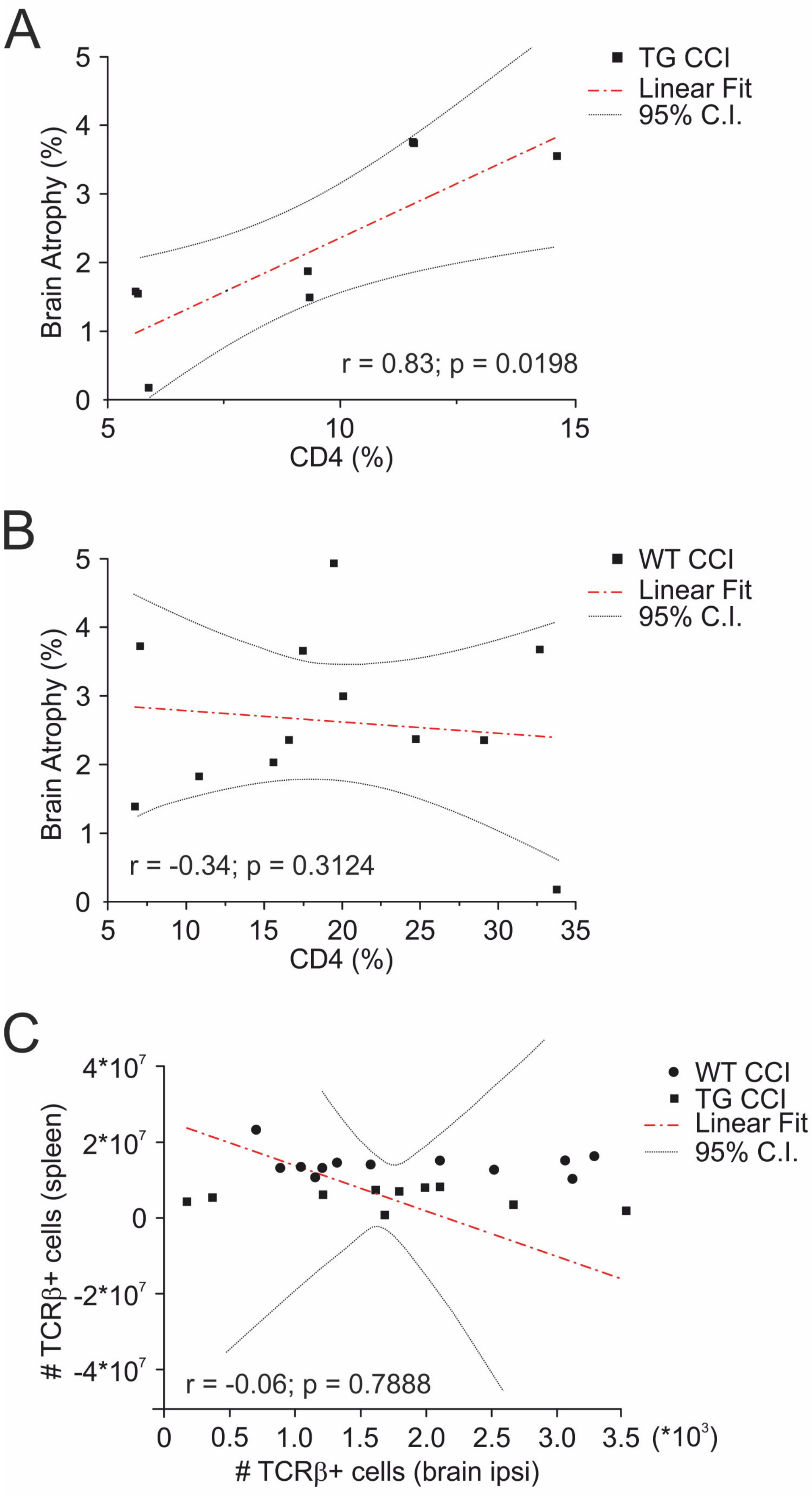


**Supplementary Figure 3:** CD4+ T cell frequency directly correlate with the percentage of tissue loss in TG CCI **(A)** but not in WT CCI mice **(B)**. Scatter plot in **(C)** show the correlation between the calculated total number of T cells in the spleen and the number of infiltrating T cells in the perilesional cortex in each analyzed mouse. Pearson's linear regression.

### Supplementary Figure 4

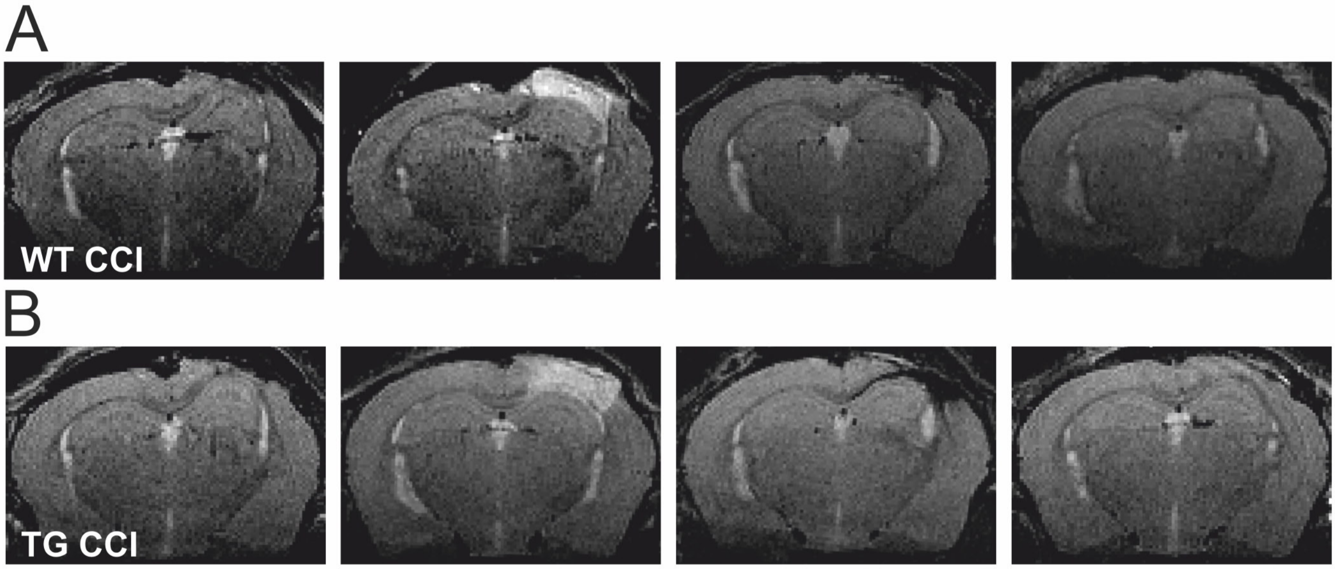


**Supplementary Figure 4:** Different types of lesions observed after CCI induction. **(A)** WT CCI mice **(B)**. TG CCI mice.

### Supplementary Table 1

| Genotype | Age (days) | Weight (g) | Anesthesia (isoflurane % during mantainance) | Surgery duration (min) | Anesthesia (isoflurane total volume administered; mL) | Impact velocity (m/s) | Injury (EDH/SDH/SAH) |
|---|---|---|---|---|---|---|---|
| **C57Bl/6JOla** | 222 ± 50 | 34.7 ± 3.4 | 1.2 ± 0.0 | 36 ± 2 | 2.04 ± 0.06 | 5.01 ± 0.01 | 4/4/4 |
| **K14-VEGFR3-Ig** | 203 ± 39 | 36.2 ± 3.3 | 1.0 ± 0.1 | 39 ± 4 | 2.10 ± 0.08 | 5.01 ± 0.01 | 1/5/5 |

**Supplementary Table 1:** ***Variables related to TBI-induction procedures.*** Standardization of procedures resulted in no statistical differences in the related variables between WT (C57Bl/6OlaJ) and TG (K14-VEGFR3-Ig) mice. Data are relative to the animals considered for the final analyses (spleen - see main text for details). The total volume of isoflurane administered during the surgery refers to the ”induction” and ”mantainance” phases. Epidural hematoma (EDH), subdural hematoma (SDH) and subarachnoid hemorrhage (SAH) were evaluated immediately after TBI induction by visual inspection of the injury area, by a trained researcher.

### Supplementary Table 2

#### BRAIN

**CD8+CD44$^{hi}$CD69+**

| | TG ipsi ChiSq | TG ipsi Mean Ranks | TG ipsi p value | WT contra ChiSq | WT contra Mean Ranks | WT contra p value |
|---|---|---|---|---|---|---|
| *WT ipsi* | 0.1263 | 10.00/9.10 | 0.7338 | --- | 4.60/9.70 | 0.040 (*) |
| *TG contra* | --- | 7.14/8.75 | 0.5400 | 0.7978 | 7.28/5.40 | 0.3973 |

**CD8+CD44$^{hi}$CD69-**

| | TG ipsi ChiSq | TG ipsi Mean Ranks | TG ipsi p value | WT contra ChiSq | WT contra Mean Ranks | WT contra p value |
|---|---|---|---|---|---|---|
| *WT ipsi* | 0.032 | 9.75/9.30 | 0.8652 | --- | 11.40/6.30 | 0.040 (*) |
| *TG contra* | --- | 8.86/7.25 | 0.5400 | 0.7978 | 5.71/7.60 | 0.3973 |

**CD8+CD44$^{neg}$CD69+**

| | TG ipsi ChiSq | TG ipsi Mean Ranks | TG ipsi p value | WT contra ChiSq | WT contra Mean Ranks | WT contra p value |
|---|---|---|---|---|---|---|
| *WT ipsi* | 0.0079 | 9.37/9.60 | 0.9324 | --- | 4.80/9.60 | 0.053 |
| *TG contra* | --- | 7.57/8.37 | 0.7800 | 2.5592 | 7.86/4.60 | 0.1123 |

**CD8+CD44$^{neg}$CD69-**

| | TG ipsi ChiSq | TG ipsi Mean Ranks | TG ipsi p value | WT contra ChiSq | WT contra Mean Ranks | WT contra p value |
|---|---|---|---|---|---|---|
| *WT ipsi* | 5.779 | 6.12/12.20 | 0.0101(**) | --- | 6.70/8.65 | 0.460 |
| *TG contra* | --- | 6.00/9.75 | 0.090 | 0.5566 | 6.00/7.20 | 0.4821 |

**CD4+CD44$^{hi}$CD69+**

| | TG ipsi ChiSq | TG ipsi Mean Ranks | TG ipsi p value | WT contra ChiSq | WT contra Mean Ranks | WT contra p value |
|---|---|---|---|---|---|---|
| *WT ipsi* | 0.109 | 10.50/9.63 | 0.7514 | --- | 7.50/9.81 | 0.400 |
| *TG contra* | --- | 8.00/8.00 | 1.000 | 1.6530 | 8.28/5.50 | 0.2118 |

**CD4+CD44$^{hi}$CD69-**

| | TG ipsi ChiSq | TG ipsi Mean Ranks | TG ipsi p value | WT contra ChiSq | WT contra Mean Ranks | WT contra p value |
|---|---|---|---|---|---|---|
| *WT ipsi* | 0.197 | 8.87/10.00 | 0.6704 | --- | 11.19/16.64 | 0.56 |
| *TG contra* | --- | 7.57/8.37 | 0.780 | 1.3061 | 5.85/8.33 | 0.271 |

**CD4+CD44$^{neg}$CD69+**

| | TG ipsi ChiSq | TG ipsi Mean Ranks | TG ipsi p value | WT contra ChiSq | WT contra Mean Ranks | WT contra p value |
|---|---|---|---|---|---|---|
| *WT ipsi* | 1.254 | 11.69/8.77 | 0.275 | --- | 8.25/9.40 | 0.680 |
| *TG contra* | --- | 5.92/9.81 | 0.094 | 0.325 | 6.50/7.58 | 0.5911 |

**CD4+CD44$^{neg}$CD69-**

| | TG ipsi ChiSq | TG ipsi Mean Ranks | TG ipsi p value | WT contra ChiSq | WT contra Mean Ranks | WT contra p value |
|---|---|---|---|---|---|---|
| *WT ipsi* | 0.441 | 11.00/9.27 | 0.5222 | --- | 11.00/7.90 | 0.240 |
| *TG contra* | --- | 8.00/8.00 | 1.000 | 0.2032 | 6.57/7.50 | 0.6718 |

**Supplementary Table 2:** Statistical analyses of the CD8+ and CD4+ T cell subpopulation frequencies was performed using the Kruskal Wallis test or the paired samples Wilcoxon signed ranked test. Bonferroni correction was used for multiple analyses (refer to main text for detailed information). * $p < 0.05$; ** $p > 0.01$ by paired samples Wilcoxon signed ranked test.

### Supplementary Table 3

| SPLEEN | ChiSq | Mean Ranks | p value | ChiSq | Mean Ranks | p value |
|---|---|---|---|---|---|---|
| **CD8+CD44$^{hi}$ CD69+** | *TG CCI* | | | *WT naïve* | | |
| *WT CCI* | 12.631 | 14.50/5.50 | 0.0004 (***) | 2.719 | 9.00/13.58 | 0.1000 |
| *TG naïve* | 0.521 | 9.94/8.17 | 0.4881 | 14.737 | 17.00/6.50 | 0.0001(***) |
| **CD8+CD44$^{hi}$ CD69-** | *TG CCI* | | | *WT naïve* | | |
| *WT CCI* | 8.347 | 13.56/6.25 | 0.0012(**) | 1.828 | 9.45/13.21 | 0.1825 |
| *TG naïve* | 1.689 | 10.69/7.50 | 0.2031 | 2.020 | 13.22/9.33 | 0.1603 |
| **CD8+CD44$^{int}$ CD69-** | *TG CCI* | | | *WT naïve* | | |
| *WT CCI* | 3.321 | 6.94/11.55 | 0.0663 | 1.045 | 9.95/12.79 | 0.3183 |
| *TG naïve* | 3.000 | 6.75/11.00 | 0.0825 | 0.323 | 1.89/10.33 | 0.5829 |
| **CD8+CD44$^{neg}$ CD69+** | *TG CCI* | | | *WT naïve* | | |
| *WT CCI* | 5.755 | 12.87/6.80 | 0.0113(*) | 0.004 | 11.40/11.58 | 0.9493 |
| *TG naïve* | 2.370 | 11.00/7.22 | 0.1271 | 1.459 | 12.89/9.58 | 0.2363 |
| **CD8+CD44$^{neg}$ CD69-** | *TG CCI* | | | *WT naïve* | | |
| *WT CCI* | 7.595 | 6.25/12.60 | 0.0024(**) | 1.655 | 13.45/9.87 | 0.2057 |
| *TG naïve* | 0.231 | 8.37/9.56 | 0.6456 | 7.293 | 6.78/14.17 | 0.0037(**) |

| | ChiSq | Mean Ranks | p value | ChiSq | Mean Ranks | p value |
|---|---|---|---|---|---|---|
| **CD4+CD44$^{hi}$ CD69+** | *TG CCI* | | | *WT naïve* | | |
| *WT CCI* | 12.008 | 14.37/5.60 | 0.0005(***) | 0.353 | 12.40/10.75 | 0.5655 |
| *TG naïve* | 0.454 | 9.87/8.22 | 0.5182 | 14.727 | 17.00/5.50 | 0.0001(***) |
| **CD4+CD44$^{hi}$ CD69-** | *TG CCI* | | | *WT naïve* | | |
| *WT CCI* | 11.400 | 14.25/5.70 | 0.0007(***) | 6.787 | 7.55/14.79 | 0.0058(**) |
| *TG naïve* | 4.083 | 6.37/11.33 | 0.0386(*) | 10.227 | 16.00/7.25 | 0.0003(***) |
| **CD4+CD44$^{int}$ CD69-** | *TG CCI* | | | *WT naïve* | | |
| *WT CCI* | 12.631 | 4.50/13.50 | 0.0003(***) | 1.332 | 13.25/10.04 | 0.2581 |
| *TG naïve* | 0.148 | 9.50/8.55 | 0.7133 | 14.727 | 5.00/15.50 | 0.0001(***) |
| **CD4+CD44$^{neg}$ CD69+** | *TG CCI* | | | *WT naïve* | | |
| *WT CCI* | 5.337 | 12.75/6.90 | 0.0156(*) | 0.480 | 10.45/12.37 | 0.5018 |
| *TG naïve* | 0.592 | 10.00/8.11 | 0.4593 | 2.913 | 13.66/9.00 | 0.0878 |
| **CD4+CD44$^{neg}$ CD69-** | *TG CCI* | | | *WT naïve* | | |
| *WT CCI* | 0.197 | 8.87/10.00 | 0.6704 | 0.000 | 11.5/11.5 | 1.0000 |
| *TG naïve* | 0.453 | 8.12/9.78 | 0.5182 | 0.409 | 10.00/11.75 | 0.5363 |

**Supplementary Table 3:** Statistical analyses of the CD8+ and CD4+ T cell subpopulation in the spleen was performed using Kruskal Wallis test with Bonferroni correction for multiple analyses (refer to main text for detailed information). * $p < 0.05$; ** $p < 0.01$; *** $p < 0.001$
